# Characterising strategy use during the performance of hippocampal-dependent tasks

**DOI:** 10.1101/807990

**Authors:** Ian A. Clark, Anna M. Monk, Eleanor A. Maguire

## Abstract

Recalling the past, thinking about the future and navigating in the world are linked with a brain structure called the hippocampus. Precisely how the hippocampus enables these critical cognitive functions is still debated. The strategies people use to perform tasks associated with these functions have been under-studied, and yet such information could augment our understanding of the associated cognitive processes and neural substrates. Here, we devised and deployed an in-depth protocol to examine the explicit strategies used by 217 participants to perform four naturalistic tasks widely acknowledged to be hippocampal-dependent, namely, those assessing scene imagination, autobiographical memory recall, future thinking and spatial navigation. In addition, we also investigated strategy use for three laboratory-based memory tasks, one of which is held to be hippocampal-dependent – concrete verbal paired associates – and two tasks which are likely hippocampal-independent – abstract verbal paired associates and the dead or alive semantic memory test. We found that scene visual imagery was the dominant strategy not only when mentally imagining scenes, but also during autobiographical memory recall, when thinking about the future and during navigation. Moreover, scene visual imagery strategies were used most frequently during the concrete verbal paired associates task, whereas verbal strategies were most prevalent for the abstract verbal paired associates task and the dead or alive semantic memory task. The ubiquity of specifically scene visual imagery use across a range of tasks may attest to its, perhaps underappreciated, importance in facilitating cognition, whilst also aligning with perspectives that emphasise a key role for the hippocampus in constructing scene imagery.

## INTRODUCTION

We recall our past experiences in the form of autobiographical memories, and are also able to imagine potential future events. Along with spatial navigation (O’Keefe and Nadel, 1978; Maguire et al., 2000; Ekstrom et al., 2003; Moser et al., 2008), these cognitive functions have been linked with a brain structure called the hippocampus (Scoville and Milner, 1957; Tulving, 1985; Klein et al., 2002; Svoboda et al., 2006; Addis et al., 2007; Verfaellie and Keane, 2017). Views differ on the precise role played by the hippocampus in supporting these seemingly disparate cognitive functions (e.g., Schacter et al., 2012; Maguire and Mullally, 2013; Moscovitch et al., 2016; Ekstrom and Ranganath, 2018).

One suggestion is that what these cognitive functions have in common is the prominent involvement of scene visual imagery (Hassabis and Maguire, 2007; Maguire and Mullally, 2013; see also, Rubin and Umanath, 2015; Robin, 2018 for related theoretical viewpoints). A scene is as a naturalistic three-dimensional, spatially coherent representation of the world typically populated by objects and viewed from an egocentric perspective (Dalton et al., 2018). The ability to construct scene visual imagery is impaired in patients with bilateral hippocampal damage (Hassabis et al., 2007a; Rosenbaum et al., 2009; Andelman et al., 2010; Race et al., 2011; Mullally et al., 2012; Maguire and Mullally, 2013), while imagination of single objects remains intact (Hassabis et al., 2007a). Similarly, neuroimaging studies have consistently reported hippocampal engagement when healthy participants imagine visual scenes in comparison to, for example, single objects (Hassabis et al., 2007b; Andrews-Hanna et al., 2010; Zeidman and Maguire, 2016; Barry et al., 2018; Clark et al., 2018; Dalton et al., 2018; Palombo et al., 2018; Robin, 2018).

A recent examination of people’s ability to imagine scenes, recall their autobiographical memories, think about the future and navigate showed that the capacity to construct scene visual imagery mediated performance across tasks assessing these functions (Clark et al., 2019). Furthermore, responses on questionnaires probing scene visual imagery were found to have a significant association with performance on scene imagination, autobiographical memory and future thinking tasks (Clark and Maguire, 2020). Although findings such as these indirectly implicate scene visual imagery across what might be regarded as ‘naturalistic’ hippocampal-dependent tasks, there is a dearth of studies directly examining the explicit strategies people actually use during their performance (Andrews-Hanna et al., 2010).

In contrast to naturalistic tasks like autobiographical memory recall, the use of cognitive strategies during simpler, laboratory-based memory tasks, such as word list learning, has been studied more extensively. In this domain, strategies have been found to differ in terms of their modality, including visual imagery and verbal strategies involving sentences or stories, and in their complexity, ranging from simple strategies like rote repetition to more complex strategies involving bizarre and distinct visual imagery and interactive visual scenes (Roberts, 1968; Paivio, 1969; Boltwood and Blick, 1970; Bower, 1970; Stoff and Eagle, 1971; McDaniel and Kearney, 1984; Kroll et al., 1986; McDaniel and Einstein, 1986; Einstein and McDaniel, 1987; Marschark and Hunt, 1989; Logie et al., 1996; Hertzog et al., 1998; Dunlosky and Kane, 2007). Within the realm of laboratory-based studies, suggestions have also been offered about methods for investigating strategy use. One recommendation is that strategy data be collected using “think aloud” protocols on a trial-by-trial basis, that is, by asking participants to report what is going through their minds as they are performing the task in question (e.g., Ericsson and Simon, 1980). Advantages of this concurrent approach include reduced forgetting of the strategies used (e.g., Dunlosky and Hertzog, 1998; Dunlosky and Hertzog, 2001), and the scope for participants to report the use of multiple strategies (e.g., Siegler, 1988).

It is difficult, however, to extrapolate this method directly to naturalistic tasks such as autobiographical memory recall and future thinking. For example, it is simply not possible to have a participant describe a memory or a future scenario out loud while simultaneously reporting the strategies they used to do so. Moreover, compared to laboratory-based tasks, it is difficult to define what constitutes a specific trial, because each scene, memory or future scenario description can last for several minutes. While one could ask about strategy use at different points within the description, this would likely disrupt the narrative flow, as well as these requests being arbitrary in their occurrence. It is also usually the case that established protocols for interrogating strategies are typically designed to elicit information about a single task. This presents a challenge when investigating strategy use across multiple tasks performed by the same participants, because probing strategies for one task could then influence how subsequent tasks are performed (Dunlosky and Hertzog, 2001).

In the current study we sought to build on the strategy use research of laboratory-based tasks to examine the explicit strategies used in naturalistic tasks that are held to be hippocampal-dependent, namely, scene imagination, autobiographical memory recall, future thinking and spatial navigation. We first developed a new protocol for strategy data collection that could be used for these tasks, and for examining strategy use across multiple tasks performed by the same participants. This resulted in a strategy use questionnaire tailored to each task of interest which could be completed by participants retrospectively after performing all of the tasks. Each questionnaire started with a reminder of the task in question to help participants think back to the task and how they performed it. A wide range of potential strategies (between 12 and 24) was then provided to the participants and they were asked to select all of the strategies they used for that task. Finally, participants provided a rank for each strategy relating to its degree of use.

We regarded this as the most parsimonious approach for several reasons. While collecting strategy information concurrently is often deemed to be preferable to probing strategy use retrospectively, in fact there are acknowledged advantages and challenges associated with each method (Dunlosky and Hertzog, 2001). Retrospective reports are an efficient methodology especially when the use of concurrent reports is not possible and, as in our case, it includes task reminders to reduce the forgetting that can accompany retrospective protocols. We were also able to assess variations in strategy use within the same task (e.g., as detailed by Siegler, 1988) by including a variety of different strategy options and encouraging participants to indicate all the strategies they used to perform each task. Furthermore, by gathering information about how much each strategy was used, we could examine which strategies were the most important for each person for every task.

We collected strategy use data from 217 participants for seven tasks. These included the four hippocampal-dependent naturalistic tasks of primary interest which assessed scene imagination, autobiographical memory recall, future thinking and spatial navigation. We also examined three laboratory-based memory tasks – concrete verbal paired associates (VPA), abstract VPA (Clark et al., 2018) and the dead or alive semantic memory task (Kapur et al., 1989). The inclusion of the latter tasks allowed us to assess whether or not any strategy use patterns we observed were simply due to the naturalistic nature of the main tasks of interest. The laboratory-based tasks also enabled us to compare strategy use between tasks known to be hippocampal-dependent, which included the concrete VPA task (Zola-Morgan et al., 1986; Squire, 1992; Spiers et al., 2001; Clark et al., 2018), and those that are held to be hippocampal-independent, namely the abstract VPA and semantic memory tasks (Binder and Desai, 2011; Clark et al., 2018). We hypothesised that scene visual imagery strategies would predominate for the hippocampal-dependent tasks (scene imagination, autobiographical memory recall, future thinking, navigation and concrete VPA), while this would not be the case the hippocampal-independent tasks (abstract VPA and the dead or alive task). These predictions align with perspectives that place the construction of scene imagery at the heart of hippocampal processing (Hassabis and Maguire, 2007; Maguire and Mullally, 2013; Clark et al., 2019).

## MATERIALS AND METHODS

### Participants

Two hundred and seventeen people took part in the study, 109 females and 108 males. They were aged between 20 and 41 years of age, had English as their first language, and reported no psychological, psychiatric, neurological or behavioural health conditions. Participants were recruited from the general population to ensure wide sampling of task performance and strategy use. The age range was restricted to 20-41 to limit any possible effects of ageing. The mean age of the sample was 29.0 years (SD = 5.60). Participants reporting hobbies or vocations known to be associated with the hippocampus (e.g., licensed London taxi drivers) were excluded. Participants were reimbursed £10 per hour for taking part which was paid at study completion. The study was approved by the University College London Research Ethics Committee. All participants gave written informed consent in accordance with the Declaration of Helsinki.

### Procedure

Participants first completed the tasks over three separate testing sessions. The order of the tasks within each visit was the same for all participants (see Clark et al., 2019). Task order was arranged so as to avoid interference, for example, not having a verbal task followed by another verbal task, and to provide sessions of approximately equal length (∼3-3.5 hours, including breaks). Strategy data were collected in a separate final session, after all the tasks had been completed.

All of the cognitive tasks that we used are published. Here, for convenience, we describe each task briefly.

### Naturalistic tasks

#### Scene construction task (Hassabis et al., 2007a)

Participants are required to mentally construct visual scenes of commonplace settings. For each scene, a short cue is provided (e.g., imagine lying on a beach in a beautiful tropical bay), and the participant is asked to imagine the scene that is evoked and then describe it out loud in as much detail as possible. Participants are explicitly told not to describe a memory, but to create a new scene that they have never experienced before.

#### Autobiographical Interview (AI; Levine et al., 2002)

Participants are asked to provide autobiographical memories from a specific time and place over four time periods – early childhood (up to 11 years of age), teenage years (from 11-17 years of age), adulthood (from 18 years of age up to 12 months prior to the interview; two memories are requested) and the last year (a memory from the last 12 months).

#### Future thinking task (Hassabis et al., 2007a)

This task follows the same procedure as the scene construction task, but requires participants to imagine three plausible future scenes involving themselves (an event at the weekend; next Christmas; the next time they meet a friend). Participants are explicitly told not to describe a memory, but to create a new future scene.

#### Navigation tasks (Woollett and Maguire, 2010)

Navigational learning is examined using movies of navigation through an unfamiliar town involving two overlapping routes, which are shown to participants four times. Five tasks are then conducted to examine how well participants learned the town. First, following each viewing of the route movies, participants are shown four short movie clips – two from the actual routes, and two distractors. Participants indicate whether they had seen each movie clip or not. Second, after all four route viewings are completed, recognition memory for scenes from the routes is tested. A third task involves assessing knowledge of the spatial relationships between landmarks from the routes in the form of proximity judgements. Fourth, route knowledge is examined by having participants place photographs from the routes in the correct order as if travelling through the town. Finally, participants draw a sketch map of the two routes including as many landmarks as they can remember.

### Laboratory-based memory tasks

#### Concrete VPA (Clark et al., 2018)

The concrete VPA is based upon the Wechsler Memory Scale IV VPA task (Wechsler, 2009). Participants are asked to learn and then remember 14 word pairs, made up of concrete, high imagery words. Learning takes place over four trials, where each time (in a different order) the 14 word pairs are read out to the participant. Following this, the first word of each pair is given and the participant is asked for the corresponding word, with feedback (i.e., the correct answer is provided if necessary). After 30 minutes, the participants are tested again in the same way but without feedback. Participants are not told about the delayed recall test in advance.

#### Abstract VPA (Clark et al., 2018)

The abstract VPA is identical to the concrete VPA with one important difference. Instead of using concrete, high imagery words, it uses only abstract, very low imagery words. Importantly, the words in the abstract VPA task are highly matched with those in the concrete VPA task in terms of linguistic characteristics (e.g., length, phonemes and syllables) and frequency of use in the English language. This allows for two very similar tasks to be assessed, where one (the concrete VPA) is thought to be hippocampal-dependent, while the other (the abstract VPA) is not (Maguire and Mullally, 2013; Clark and Maguire, 2016; Clark et al., 2018).

#### Dead or alive task (Kapur et al., 1989)

This is a test of semantic knowledge. Participants are presented with the names of 74 famous individuals and are first asked to remove any names that they do not recognise. For those that the participant knows, they are then asked to indicate whether the individual is dead or alive.

#### Strategy use

There is currently no standard methodology for studying strategy use in the context of these cognitive tasks. We therefore designed a novel protocol for collecting and analysing detailed strategy information for each cognitive task.

#### Identification of strategies

To identify possible strategies used to perform the tasks, 30 participants were recruited who did not take part in the main study (15 female; mean age 27.07 years, SD = 7.32). Participant recruitment was based on an individual’s general use of visual imagery. The use of visual imagery is a well-known strategy (Paivio, 1969; Andrews-Hanna et al., 2014; Greenberg and Knowlton, 2014) and we wanted to represent all types of strategies, not just those that are based on visual imagery. General visual imagery use was determined via the Spontaneous Use of Imagery Scale (SUIS; Reisberg et al., 2003), where scores can range from 12 (very low/no spontaneous use of visual imagery) to 60 (high spontaneous use of visual imagery). Fifteen participants were recruited who reported high scores on the SUIS (≥ 46) and 15 with low SUIS scores (≤ 40), with 11 of these low scoring participants reporting values ≤ 31. The average score of the participants in this identification of strategies study was 40.03 (SD = 9.97) with a range from 24 to 57.

To collect information on individual task strategies, participants first performed the cognitive tasks, after which they were asked open-ended questions about the strategies they employed for each task. Participants were encouraged to report all strategies that they used for a task in as much detail as possible, regardless of how much or little they used them.

Strategy responses from the participants were then combined with any relevant additional strategies identified from the extant literature. Examination of all of these strategies highlighted both specific and more general techniques that participants used to perform each of the tasks. From this, we generated a large number of strategy statements, ranging from 12 to 24 strategies, for each task.

The strategies generated for each task are provided in the Supplementary Material.

#### Strategy questionnaires

The information from the strategy identification study was used to construct a strategy questionnaire for each task for use with the participants in the main experiment. The questionnaires were presented on a computer screen and were participant-paced and led, but with the involvement of the experimenter where required. The strategy questionnaires were administered in a separate session after all the tasks had been completed. This session focused solely on collecting the strategy information for each task. The average length of time between the final testing session and the strategy questionnaires session was 6.05 days (SD = 5.33).

Three steps were involved in data collection. First, a brief reminder of the task was presented. Second, participants selected the strategies they used for that task from the extensive list of possible strategies. Third, participants ranked their selected strategies in relation to their degree of use.

Strategies were requested for scene construction, each memory age of the AI, future thinking, for the learning of the town in the navigation task and each of the five navigation tasks, for the learning and delayed recall of the concrete and also the abstract VPA tasks, and for the dead or alive task.

##### Task reminder

The task reminder varied according to the task. For some tasks, a picture of the task was presented, while for others the tasks were verbally described. The experimenter then ensured that the participant fully remembered the task (providing additional information if required) before the participant moved on to the strategy selection.

##### Strategy selection

Following the task reminder, all the possible strategies for the task that were generated from the strategy identification study were presented as a list on a computer screen. For each strategy, participants were requested to respond either “Yes” (that they used the strategy) or “No” (that they did not use the strategy). A response was required for every strategy to ensure that none were accidently overlooked. It was made clear that selecting one strategy did not preclude the selection of any of the others, as more than one strategy could be deployed during a task.

##### “Other” strategies

While our strategy list for each task was extensive, we also accounted for the possibility that participants may have used strategies that were not included on the list. As such, for all tasks the option “Other”, with space to describe new strategies, was also available.

##### Strategy ranking

A list of the strategies that a participant indicated they used during the task was then presented to them. They were asked to rank each of the strategies according to how much of the time they used them. Outside of these instructions they were free to indicate any form of ranking. Thus, if they felt they used multiple strategies equally this could be indicated. For example, if three strategies were chosen they could be ranked:

- 1, 2, 3 – where the strategy ranked 1 was used most of the time, followed by the strategy ranked 2, and then the strategy ranked 3.
- 1, 1, 1 – where all strategies were used equally.
- 1, 2, 2 – where one strategy was used the most, and the other two less frequently, but the secondary strategies were used equally.

##### Question order

The task reminders and strategy selection were presented in two orders (with half the participants doing each order) to reduce the possibility of order effects. For order 1 the task order was: concrete VPA, AI, abstract VPA, navigation, dead or alive, scene construction, future thinking. The strategies were listed with visual imagery strategies first, followed by verbal strategies. For order 2, the task order was reversed, and the strategies were listed starting with verbal strategies first followed by visual imagery strategies.

## Data analysis

For the AI, strategies were examined for each memory age separately and were then combined across all four memory ages to provide an overall autobiographical memory recall strategy. For navigation, strategies were examined separately for the learning phase during movie viewing, and the five navigation tasks, and then by combining across the five navigation tasks to provide an overall navigation strategy. For the concrete VPA, and also the abstract VPA, strategies were examined for the learning and delayed recall phases separately.

For all tasks, we focused specifically on rank 1 strategies, that is, the strategy or strategies that the participant used most often and deemed the most important for that task.

If “Other” responses were provided, these were examined to ascertain if the description closely resembled a strategy that was already listed and, if this was the case, the strategy was reallocated from Other to that strategy. There was no situation where the Other description referred to a new strategy that was not already represented on the list (a detailed breakdown of Other responses and their reallocations is provided in the Supplementary Material).

To streamline the data analysis process, each strategy was allocated to one of three primary strategy categories: scene visual imagery strategies, other visual imagery strategies and verbal strategies (see the Supplementary Material for all of the strategies generated for each task, and into which primary category they were placed). Note that participants were not aware of the strategy category distinctions. In general, a scene visual imagery strategy was one which evoked a visual image of a scene, that is, the visual imagery had a sense of depth and background. Other visual imagery strategies evoked visual imagery, but this could not be defined as a scene. There was no sense of depth or background, a typical example being a visual image of a single object. A verbal strategy was one which evoked no visual imagery at all, with reliance instead upon words and phrases.

To compare the frequency of the primary strategies, Chi Square tests (thresholded at p <0.05) were performed across the three categories; scene visual imagery vs. other visual imagery vs. verbal. Note that all of the rank 1 choices of all the participants are reflected in the analyses and figures; thus if a participant identified multiple strategies as rank 1, all were included.

## RESULTS

### Naturalistic tasks

For the scene construction task, perhaps unsurprisingly, there was widespread use of scene visual imagery strategies compared to the other strategy types (Figure 1A; *scene visual imagery strategies =* 82.06%; *other visual imagery strategies =* 8.64%; *verbal strategies =* 9.30%; χ^2^ (2) = 321.62, p < 0.001).

**FIGURE 1.**
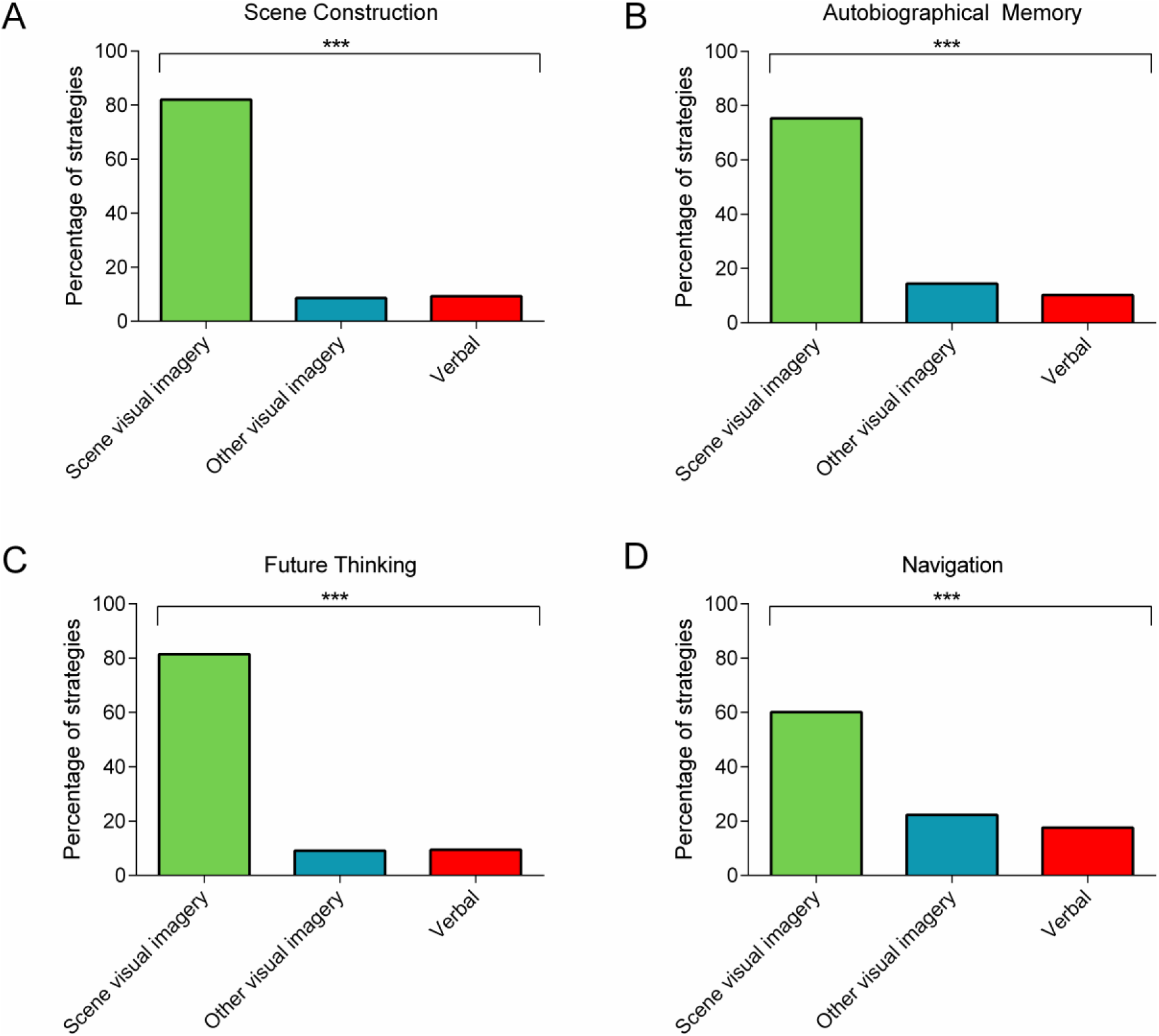
The percentage of rank 1 scene visual imagery, other visual imagery or verbal strategies used during the four naturalistic tasks involving: **(A)** scene construction; **(B)** autobiographical memory (combined across memory ages); **(C)** future thinking; **(D)** navigation (combined across the navigation tasks). Chi Square tests comparing the frequency of the rank 1 strategies across the three categories revealed a consistent use of scene visual imagery strategies compared to the other strategy types for all four tasks. There are no error bars because the graphs represent the percentage count for each strategy category. *** p < 0.001.

For autobiographical memory recall, there was also consistent use of scene visual imagery strategies compared to the other two strategy types. This was evident for all memory ages and when the data were combined across the memories (childhood: *scene visual imagery strategies =* 79.68%; *other visual imagery strategies =* 12.38%; *verbal strategies =* 7.94%; χ^2^ (2) = 305.45, p < 0.001; teenage years: *scene visual imagery strategies =* 73.83%; *other visual imagery strategies =* 15.89%; *verbal strategies =* 10.28%; χ^2^ (2) = 238.43, p < 0.001; adulthood: *scene visual imagery strategies =* 71.92%; *other visual imagery strategies =* 15.46%; *verbal strategies =* 12.62%; χ^2^ (2) = 212.83, p < 0.001; last year: *scene visual imagery strategies =* 76.0%; *other visual imagery strategies =* 14.15%; *verbal strategies =* 9.85%; χ^2^ (2) = 267.15, p < 0.001; combined (Figure 1B): *scene visual imagery strategies =* 75.35%; *other visual imagery strategies =* 14.48%; *verbal strategies =* 10.17%; χ^2^ (2) = 1018.93, p < 0.001).

For the future thinking task, scene visual imagery strategies also dominated (Figure 1C; *scene visual imagery strategies =* 81.48%; *other visual imagery strategies =* 9.09%; *verbal strategies =* 9.43%; χ^2^ (2) = 309.84, p < 0.001).

For navigation, there was a distinction between the strategies deployed during learning compared to the navigation tests. Viewing the movies while learning evoked approximately equal use of all the strategy categories (*scene visual imagery strategies =* 36.86%; *other visual imagery strategies =* 30.30%; *verbal strategies* = 32.84%; χ^2^ (2) = 3.11, p = 0.21). By contrast, performance on the post-learning tests was associated with a greater use of scene visual imagery compared to the other strategy types. This was observed when investigating each of the five navigation tasks individually, and when the data were combined across the five tasks (movie clip recognition: *scene visual imagery strategies =* 57.56%; *other visual imagery strategies =* 28.67%; *verbal strategies* = 13.77%; χ^2^ (2) = 131.77, p < 0.001; scene recognition: *scene visual imagery strategies =* 62.88%; *other visual imagery strategies =* 26.98%; *verbal strategies* = 10.15%; χ^2^ (2) = 175.79, p < 0.001; proximity judgements: *scene visual imagery strategies =* 63.04%; *other visual imagery strategies =* 14.95%; *verbal strategies* = 22.01%; χ^2^ (2) = 148.93, p < 0.001; route knowledge: *scene visual imagery strategies =* 61.01%; *other visual imagery strategies =* 17.51%; *verbal strategies* = 21.49%; χ^2^ (2) = 130.83, p < 0.001; sketch map: *scene visual imagery strategies =* 56.94%; *other visual imagery strategies =* 21.76%; *verbal strategies* = 21.30%; χ^2^ (2) = 108.39, p < 0.001; combined (Figure 1D): *scene visual imagery strategies =* 60.13%; *other visual imagery strategies =* 22.28%; *verbal strategies* = 17.59%; χ^2^ (2) = 660.62, p < 0.001).

### Laboratory-based memory tasks

For the concrete VPA, scene visual imagery strategies were more apparent at both learning and delayed recall compared to the other strategy types (learning: *scene visual imagery strategies =* 50.15%; *other visual imagery strategies =* 24.93%; *verbal strategies =* 24.93%; χ^2^ (2) = 43.38, p < 0.001; delayed recall (Figure 2A): *scene visual imagery strategies =* 44.82%; *other visual imagery strategies =* 29.97%; *verbal strategies =* 25.21%; χ^2^ (2) = 22.40, p < 0.001).

**FIGURE 2.**
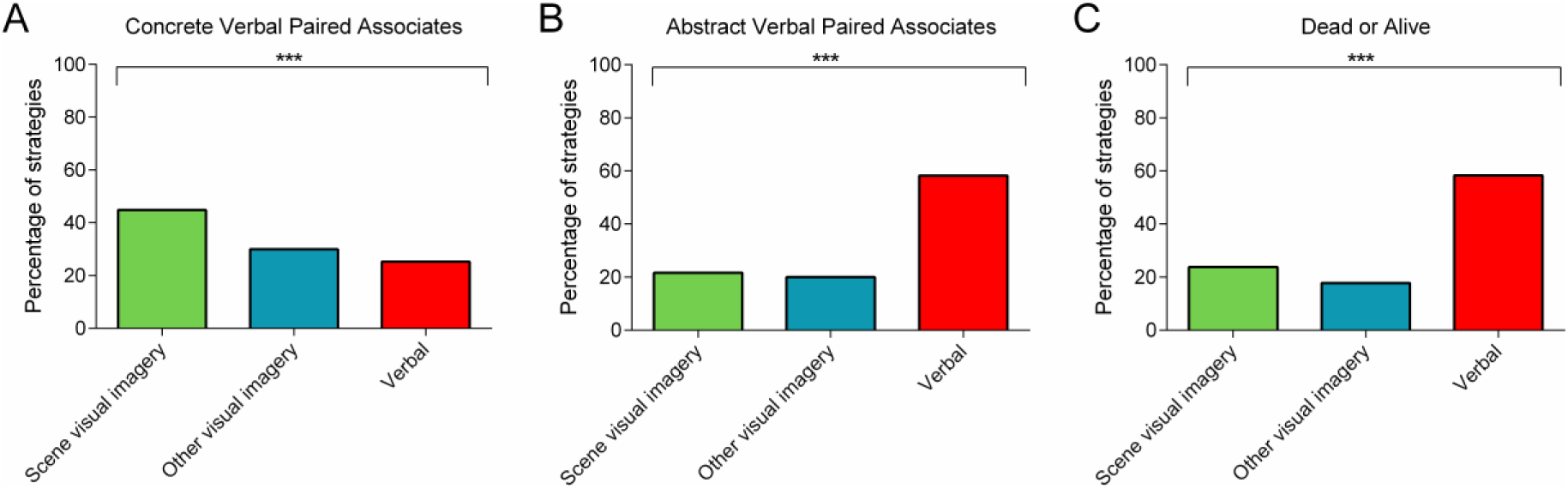
The percentage of rank 1 scene visual imagery, other visual imagery or verbal strategies used during the three laboratory-based tasks involving: **(A)** concrete verbal paired associates (delayed recall); **(B)** abstract verbal paired associates (delayed recall); **(C)** the dead or alive task. Chi Square tests comparing the frequency of the rank 1 strategies across the three categories revealed the predominant use of scene visual imagery strategies compared to the other strategy types for the concrete verbal paired associates, whereas verbal strategies dominated for the abstract verbal paired associates and the dead or alive task. There are no error bars because the graphs represent the percentage count for each strategy category. *** p < 0.001.

By contrast, and in line with our expectations, the abstract VPA elicited greater use of verbal strategies during both learning and delayed recall (learning: *scene visual imagery strategies =* 20.39%; *other visual imagery strategies =* 21.76%; *verbal strategies =* 57.85%; χ^2^ (2) = 98.30, p < 0.001; delayed recall (Figure 2B): *scene visual imagery strategies =* 21.73%; *other visual imagery strategies =* 20.06%; *verbal strategies =* 58.22%; χ^2^ (2) = 100.18, p < 0.001).

In a similar vein, verbal strategies were predominant for the dead or alive semantic memory task (Figure 2C: *scene visual imagery strategies =* 23.89%; *other visual imagery strategies =* 17.78%; *verbal strategies =* 58.33%; χ^2^ (2) = 103.27, p < 0.001).

## DISCUSSION

In this study, which involved a large sample of participants, we sought to systematically characterise the explicit strategies people used when performing a range of cognitive tasks. We found that scene visual imagery was the dominant strategy not only when mentally imagining scenes, but also during autobiographical memory recall, when thinking about the future and during navigation – all naturalistic tasks associated with the hippocampus. In addition, examination of three laboratory-based memory tasks showed that the use of scene visual imagery strategies was not limited to naturalistic tasks, whilst also indicating that participants did not invariably favour such strategies. Scene visual imagery strategies were used during concrete VPA, a task also linked to the hippocampus, whereas verbal strategies were most prevalent for tasks thought to be hippocampal-independent, namely abstract VPA and the dead or alive semantic memory task.

Behavioural experiments in healthy people have shown that the ability to construct scene visual imagery explained performance across tasks assessing autobiographical recall, future thinking and navigation (Clark et al., 2019; see also Clark and Maguire, 2020). Neuroimaging studies have also documented hippocampal engagement during the construction of scene imagery (Hassabis et al., 2007b; Zeidman and Maguire, 2016; Barry et al., 2018; Clark et al., 2018; Dalton et al., 2018). Of particular pertinence, the ability to construct scene visual imagery is impaired in patients with bilateral hippocampal damage (Hassabis et al., 2007a; Rosenbaum et al., 2009; Andelman et al., 2010; Race et al., 2011; Mullally et al., 2012; Maguire and Mullally, 2013). On the basis of this convergent evidence, the scene construction theory was proposed which suggests that the construction of scene imagery is a central function of the hippocampus, and that any task typically requiring scene imagery will be hippocampal-dependent as a consequence (Hassabis and Maguire, 2007; Maguire and Mullally, 2013). However, until now there was very little direct evidence that people actually use specifically scene visual imagery during cognitive functions such as autobiographical recall, future thinking and navigation (Andrews-Hanna et al., 2010). The current findings provide strong evidence that scene visual imagery is indeed the principal strategy deployed in such circumstances, aligning with the scene construction theory.

Why might scene imagery be used so pervasively? The most obvious answer is that it mirrors how people experience and perceive the world. In pragmatic terms it makes sense that the neural hardware and software underpinning perception would also be utilised for mental representations and during recall. In addition, scenes are a highly efficient means of packaging information. Its apparent prevalence has even led to the suggestion that scene imagery may be the currency of cognition (Maguire and Mullally, 2013).

The ubiquity of scene visual imagery affected another issue, namely, whether using such strategies actually confers any benefit for task performance. We were keen address this question. However, given the majority of participants used scene visual imagery strategies to perform the naturalistic tasks of primary interest, it was not possible to conduct meaningful analyses. For example, 212 of the 217 participants reported scene visual imagery as a rank 1 strategy during autobiographical memory recall, meaning a comparison with the remaining 5 participants would simply not have been valid.

Our results also highlight the potential perils of making assumptions about task modality. For example, as VPA tasks involve the learning and later recall of word pairs, they are typically regarded as verbal tasks. However, we found that when the words in the VPA task were imageable and concrete in nature, scene imagery strategies were used most frequently. By contrast, when the VPA task comprised low imagery abstract words, verbal strategies dominated. Therefore, focusing on the task stimuli to define task modality may not accurately reflect how participants perform a task, and underline that it is essential that task modality is assessed and not assumed.

In addition, by examining the strategies used to perform each task, we were able to make a distinction within the visual imagery domain between scene visual imagery and other visual imagery strategies. This would be difficult to assess by focusing just on task modality. Importantly, across the hippocampal-dependent tasks it was scene visual imagery strategies that were predominant. If the strategies simply reflected a visual versus verbal modality split, then a higher frequency of other visual imagery strategies might have been expected, but this is not what we found.

How would analysing strategies, instead of using assumed task modality, potentially change conclusions previously drawn? Understanding the strategies used to perform a task may, for example, allow us to better explain the relationship between what are traditionally described as verbal memory tasks and the hippocampus. The use of scene visual imagery strategies may be why concrete VPA, for example, activates the hippocampus during neuroimaging (Clark et al., 2018) and why patients with hippocampal damage, who are impaired at imagining scenes, perform so poorly on this test (e.g., Zola-Morgan et al., 1986; Spiers et al., 2001; Giovanello et al., 2003). This stands in stark contrast to the abstract VPA task comprising low imagery words which, along with the semantic memory dead or alive task, involved the use of verbal strategies. Abstract VPA stimuli do not seem to engage the hippocampus during neuroimaging (Clark et al., 2018) and, in this context, the prediction is that hippocampal-damaged patients would be relatively unimpaired on an abstract VPA task. However, the VPA tasks typically used with such patients have been concrete in nature (reviewed in Clark and Maguire, 2016), and this hypothesis remain untested. Nevertheless, our strategy data reveal a clear distinction between concrete and abstract VPA that accords with hippocampal-dependence and hippocampal-independence respectively, a distinction that is not apparent when assuming all VPA tasks are verbal.

Could it be that the strategy use protocol we employed was biased in some way towards scene visual imagery? We think this is unlikely for a number of reasons. First, a wide variety of strategies were provided, including different modalities and complexities, to avoid biasing individuals to selecting any specific strategy (see the Supplementary Materials for the full list of strategies for each task). Second, participants were presented with a list of possible strategies, and were unaware of the strategy categorisation. Third, similar numbers of potential strategies were included for each strategy category to avoid any biases towards specific categories. Fourth, the strategies were presented in two orders, with half the participants seeing the visual imagery strategies first and half the verbal strategies first, thus reducing any effects of presentation order. Fifth, participants were encouraged to select any and all strategies that they used, even if the strategies seemed to contradict each other (as different strategies could be used at different times during a task). Finally, the data collection was participant-paced and led in order to avoid any potential influence of the experimenter and, overall, the experimenter was minimally involved, as the strategy use questionnaires were completed on a computer.

To investigate strategy use in naturalistic tasks like autobiographical memory and future thinking, we had to innovate beyond methods previously validated for the collection of strategy data in laboratory-based tasks (e.g., Ericsson and Simon, 1980; Dunlosky and Hertzog, 2001). This imposed some necessary constraints. By using retrospective instead of concurrent reporting, participants could have forgotten the strategies they used to perform the tasks. However, previous reports have demonstrated the effectiveness of retrospective strategy reporting (e.g., Dunlosky and Hertzog, 2001), and we provided detailed reminders to help participants remember the tasks, with no participant indicating they were unable to recall the concomitant strategies they used. By not using trial-by-trial assessments we may have missed nuances or changes in strategy use throughout the tasks. However, our participants were provided with numerous possible strategies for each task, and were encouraged to indicate all the different strategies they used during task performance, with additional information probing the extent to which each strategy was used. We were, therefore, able to collect comprehensive information about the use of multiple strategies by an individual participant for the same task. In common with concurrent reporting methods, we investigated only those strategies that could be explicitly described by participants. We acknowledge that implicit strategies also play a role in task performance, but remain challenging to discover.

We collected strategy information by having participants choose from a list of provided strategies, instead of asking them to freely describe the strategies they used, for a number of reasons. First, this ensured that all participants underwent exactly the same procedure for strategy data collection. By contrast, in a free description situation, differing levels of experimenter involvement would likely be required depending on the participant (e.g., variation in the extent of probing), which could affect the data obtained. Second, we wanted participants to consider in-depth how they performed each of the tasks in question. Providing a large range of strategy options with responses required for each option meant that all participants had to consider their use of strategies from across different modalities. Free descriptions, on the other hand, do not necessarily encourage this, and again there could be wide variability in terms of the range of options that each participant considers. Finally, providing a list of strategy options allowed us to include specific nuances within the strategies, for example, asking whether a visual image came to mind immediately or whether the image took time to form. Obtaining this information from free descriptions would be much more difficult and likely involve substantial questioning and involvement of the experimenter – something we were keen to avoid in order to reduce any potential experimenter influence. The option “Other” was also available for all tasks, where participants were able to indicate any additional strategies they felt were not represented by the lists provided. However, Other descriptions were rarely provided, and there was no situation where the Other description referred to new strategies that were not already represented on the lists, suggesting our technique did not omit any key strategies. It will be important in future studies to investigate if rates of reported strategy use are related to the methodology used to collect them.

We have alluded throughout to cognitive task-hippocampus relationships without measuring the hippocampus itself. We felt able to do this because of the many previous neuropsychological and neuroimaging findings associating scene imagination, autobiographical memory, future thinking and navigation tasks with the hippocampus. Moreover, understanding the strategies used to perform these tasks is not reliant upon hippocampal measurement. However, establishing a direct link between strategy use and the hippocampus will be an important next step. These relationships could be assessed using resting state fMRI, task-based fMRI and structural brain measurements of the hippocampus and its connectivity.

In conclusion, the strategies used to perform naturalistic tasks have been under-studied, and yet such information could augment our understanding of the associated cognitive processes and neural substrates. In a large sample of participants we identified scene visual imagery as a dominant strategy specifically in tasks associated with the hippocampus, aligning with perspectives that emphasise a link between scene processing and this brain structure.

## DATA AVAILABILITY

The test materials are all available in the published literature. The data will be made freely available once the construction of a dedicated data-sharing portal has been completed. In the meantime, the raw data supporting the conclusions of this manuscript will be made available by the authors to any qualified researcher upon request. Requests for the data can be sent to.

## ETHICS STATEMENT

This study involving human participants was approved by the University College London Research Ethics Committee (project ID: 6743/001). All participants gave written informed consent in accordance with the Declaration of Helsinki.

## AUTHOR CONTRIBUTIONS

IC and EM designed the overall study. All authors were involved in designing the strategy questionnaires. AM contributed to data collection and scoring. IC performed the statistical analyses. IC and EM wrote the first draft of the manuscript. All authors contributed to manuscript revision, read and approved the submitted version.

## FUNDING

This work was supported by a Wellcome Principal Research Fellowship to EM (101759/Z/13/Z) and the Centre by a Centre Award from Wellcome (203147/Z/16/Z).

### ACKNOWELDGEMENTS

We thank Victoria Hotchin, Gloria Pizzamiglio and Alice Liefgreen for assistance with data collection and scoring, and Narinder Kapur for providing the dead or alive test materials.

## SUPPLEMENTAL MATERIAL

Supplemental Material is available for this article.

## Supplementary Material

### Strategies for each task

The specific strategies presented to the participants for each task are shown below. Note that for each task the participants were presented only with a list of the strategies and were not aware of the strategy categories. The strategy category information is included to show how the strategies were allocated into the three primary categories for data analysis.

Participants were asked to indicate the use of each strategy by “Yes” or “No”. They could select as many strategies as were relevant to how they performed a task. Half of the participants saw the visual imagery strategies first, the other half saw the verbal strategies first.

For all tasks, the option “Other”, with space to describe new strategies, was also available.

### Scene construction task

| Strategy category |  |
| --- | --- |
| Scene visual imagery | I had an immediate visual image of the scene in my mind, with details appearing all at once as a coherent whole. It almost felt like I was really there. |
|  | I had a visual image of the scene which I imagined in a piece-by-piece manner, adding details to gradually form a coherent whole. |
|  | I had a static 2D visual image of the scene in my mind, like looking at a photograph. |
|  | When describing part of the scene to the experimenter, my visual focus was only on that specific section and I did not visualise the rest of the scene around it. |
|  | I had a very vague or fleeting sense of a visual image in my mind, which was really unclear and hazy – it was more the idea of an image in my mind than actually seeing an image itself clearly. My very vague impression is that the image was scene-like (in that I had a sense of a space or context, albeit very vague). |
| Other visual imagery | I had separate visual images of individual objects I would typically expect to find in the scene, but did not picture the scene as a coherent whole with all the objects together. |
|  | I visualised words for possible objects that I would expect to find in the scene as though written in the air. |
|  | I visualised sentences that describe the scene, as though written in the air. |
|  | I had a very vague or fleeting sense of a visual image in my mind, which was really unclear and hazy – it was more the idea of an image in my mind than actually seeing an image itself clearly. My very vague impression is that the image (choose one):<br>- had multiple elements but was not a scene<br>- involved single isolated objects<br>- I cannot describe the image |
| Verbal | I verbally thought about the scene overall, in words or sentences alone, so that the description of the scene I gave was a continuous narrative describing the scene as a coherent whole. I did this without involving visual imagery. |
|  | I verbally thought about the scene in sections, using words or sentences alone, so that I built up an overall description gradually piece-by-piece. There was no visual imagery involved in doing this. |
|  | I verbally listed, using words and sentences alone, individual objects that I would expect to typically find in the scene, without combining them into a whole. There was no visual imagery involved in doing this. |
| Other | Please describe here other strategies you used that are not listed above. |

### Autobiographical Interview (for all memory ages)

| Strategy category |  |
| --- | --- |
| Scene visual imagery | I had an immediate, fully-formed visual image of a scene or scenes from my memory that was quite immersive, like a movie playing out. |
|  | I had an immediate, fully-formed visual image of my memory, with details appearing all at once as a coherent scene, as though looking at a photograph. |
|  | I reconstructed a visual image of the memory, by adding details in a piece-by-piece manner to gradually form a coherent whole scene. |
|  | I had a very vague or fleeting sense of a visual image in my mind, which was really unclear and hazy – it was more the idea of an image in my mind than actually seeing an image itself clearly. My very vague impression is that the image was scene-like (in that I had a sense of a space or context, albeit very vague). |
| Other visual imagery | I had separate visual images of various individual features and/or people involved in the memory in isolation, but did not picture them all together within a scene or the wider context of the overall memory. |
|  | I focused on one image of a particular feature/person involved in the memory in isolation, and described my memory using that as a reference. |
|  | I imagined written in my mind the dates/numbers/short facts in regards to the memory. |
|  | I imagined written in my mind sentences that described the memory. |
|  | I had a very vague or fleeting sense of a visual image in my mind, which was really unclear and hazy – it was more the idea of an image in my mind than actually seeing an image itself clearly. My very vague impression is that the image (choose one):<br>- had multiple elements but was not a scene<br>- involved single isolated objects<br>- I cannot describe the image |
| Verbal | I verbally described, using words or sentences alone, the whole memory as a coherent story, without using visual imagery. |
|  | I verbally described, using words or sentences alone, each main element of the memory, and combined the individual elements to form a coherent story without using visual imagery. |
|  | I verbally listed, using words or sentences alone, single facts about the memory, although not in a coherent or chronological order. |
|  | I thought of the word for a single element, fact or detail in particular that stood out. |
| Other | Please describe here other strategies you used that are not listed above. |

### Future thinking task

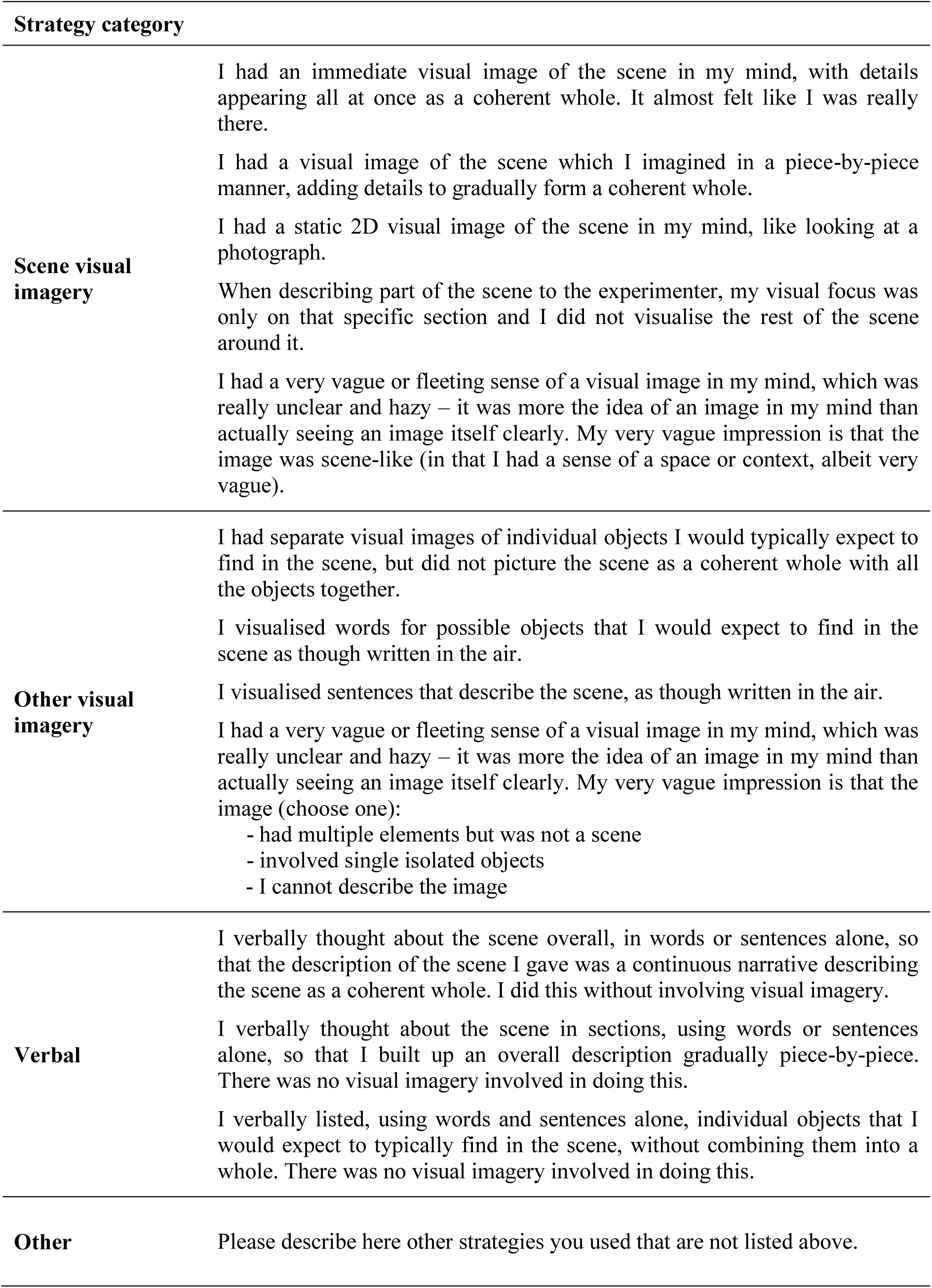

### Navigation learning

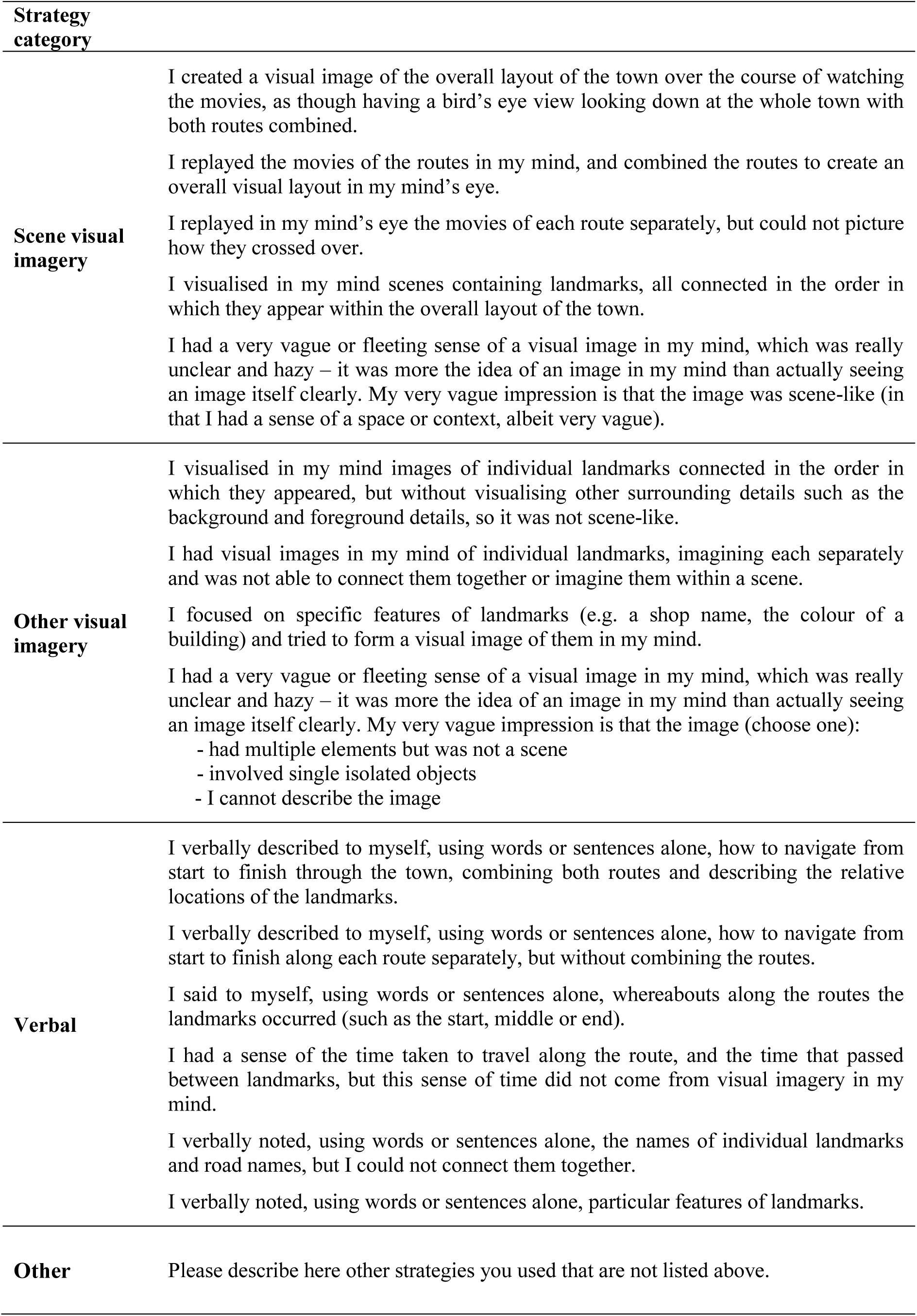

### Navigation movie clip recognition

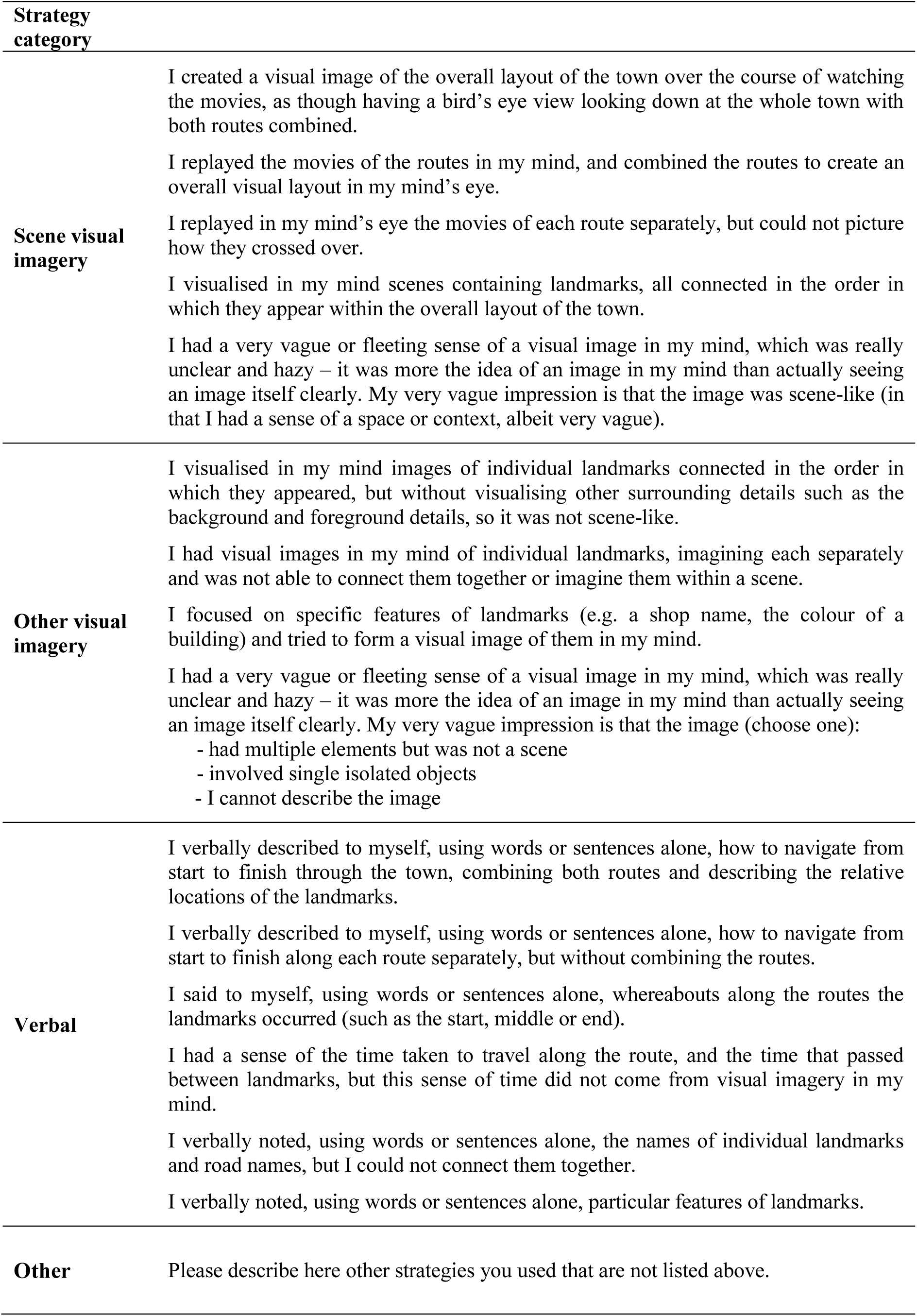

| Strategy category |  |
| --- | --- |
| **Scene visual imagery** | I created a visual image of the overall layout of the town over the course of watching the movies, as though having a bird's eye view looking down at the whole town with both routes combined.<br>I replayed the movies of the routes in my mind, and combined the routes to create an overall visual layout in my mind's eye.<br>I replayed in my mind's eye the movies of each route separately, but could not picture how they crossed over.<br>I visualised in my mind scenes containing landmarks, all connected in the order in which they appear within the overall layout of the town.<br>I had a very vague or fleeting sense of a visual image in my mind, which was really unclear and hazy – it was more the idea of an image in my mind than actually seeing an image itself clearly. My very vague impression is that the image was scene-like (in that I had a sense of a space or context, albeit very vague). |
| **Other visual imagery** | I visualised in my mind images of individual landmarks connected in the order in which they appeared, but without visualising other surrounding details such as the background and foreground details, so it was not scene-like.<br>I had visual images in my mind of individual landmarks, imagining each separately and was not able to connect them together or imagine them within a scene.<br>I focused on specific features of landmarks (e.g. a shop name, the colour of a building) and tried to form a visual image of them in my mind.<br>I had a very vague or fleeting sense of a visual image in my mind, which was really unclear and hazy – it was more the idea of an image in my mind than actually seeing an image itself clearly. My very vague impression is that the image (choose one):<br>- had multiple elements but was not a scene<br>- involved single isolated objects<br>- I cannot describe the image |
| **Verbal** | I verbally described to myself, using words or sentences alone, how to navigate from start to finish through the town, combining both routes and describing the relative locations of the landmarks.<br>I verbally described to myself, using words or sentences alone, how to navigate from start to finish along each route separately, but without combining the routes.<br>I said to myself, using words or sentences alone, whereabouts along the routes the landmarks occurred (such as the start, middle or end).<br>I had a sense of the time taken to travel along the route, and the time that passed between landmarks, but this sense of time did not come from visual imagery in my mind.<br>I verbally noted, using words or sentences alone, the names of individual landmarks and road names, but I could not connect them together.<br>I verbally noted, using words or sentences alone, particular features of landmarks. |
| **Other** | Please describe here other strategies you used that are not listed above. |

### Navigation scene recognition

| Strategy category |  |
| --- | --- |
| **Scene visual imagery** | I visually replayed parts of the route in my mind to see if the scene on the screen matched any of my mental images of the films.<br>I had visual mental snapshots of scenes from the routes, in order to see if the scene on the screen matched any of my mental images.<br>I imagined the movies being replayed on the computer screen, so that the visual image in my mind was of seeing the screen with the movie playing. This helped me to see if I recognised each scene.<br>I had a very vague or fleeting sense of a visual image in my mind, which was really unclear and hazy – it was more the idea of an image in my mind than actually seeing an image itself clearly. My very vague impression is that the image was scene-like (in that I had a sense of a space or context, albeit very vague). |
| **Other visual imagery** | I had visual images in my mind of individual, isolated landmarks, without visualising other surrounding details such as the background and foreground details, to see if any matched the scene.<br>I pictured in my mind specific features of isolated landmarks (e.g. a shop name, the colour of a building) to see if they matched the scene.<br>I had a very vague or fleeting sense of a visual image in my mind, which was really unclear and hazy – it was more the idea of an image in my mind than actually seeing an image itself clearly. My very vague impression is that the image (choose one):<br>- had multiple elements but was not a scene<br>- involved single isolated objects<br>- I cannot describe the image |
| **Verbal** | I verbally described the route to myself, using words or sentences alone, from start to finish to see if the scene matched any point on the route description.<br>I verbally thought, using words and sentences alone, of the names of individual isolated landmarks and road names, to see if they were in the scene on the screen.<br>I verbally recalled, using words or sentences alone, particular features of landmarks to see if they were in the scene on the screen. |
| **Other** | Please describe here other strategies you used that are not listed above. |

### Navigation proximity judgements

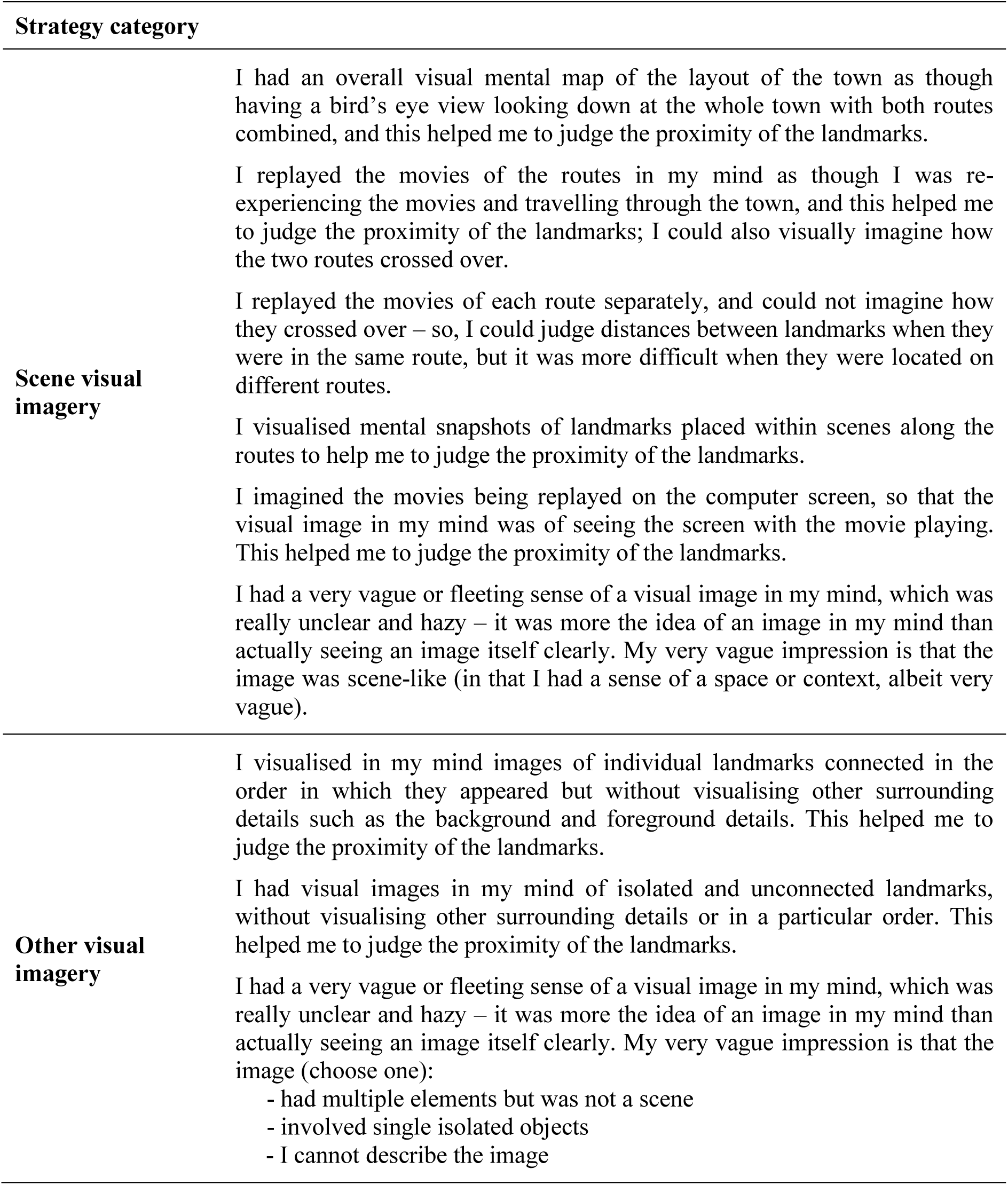

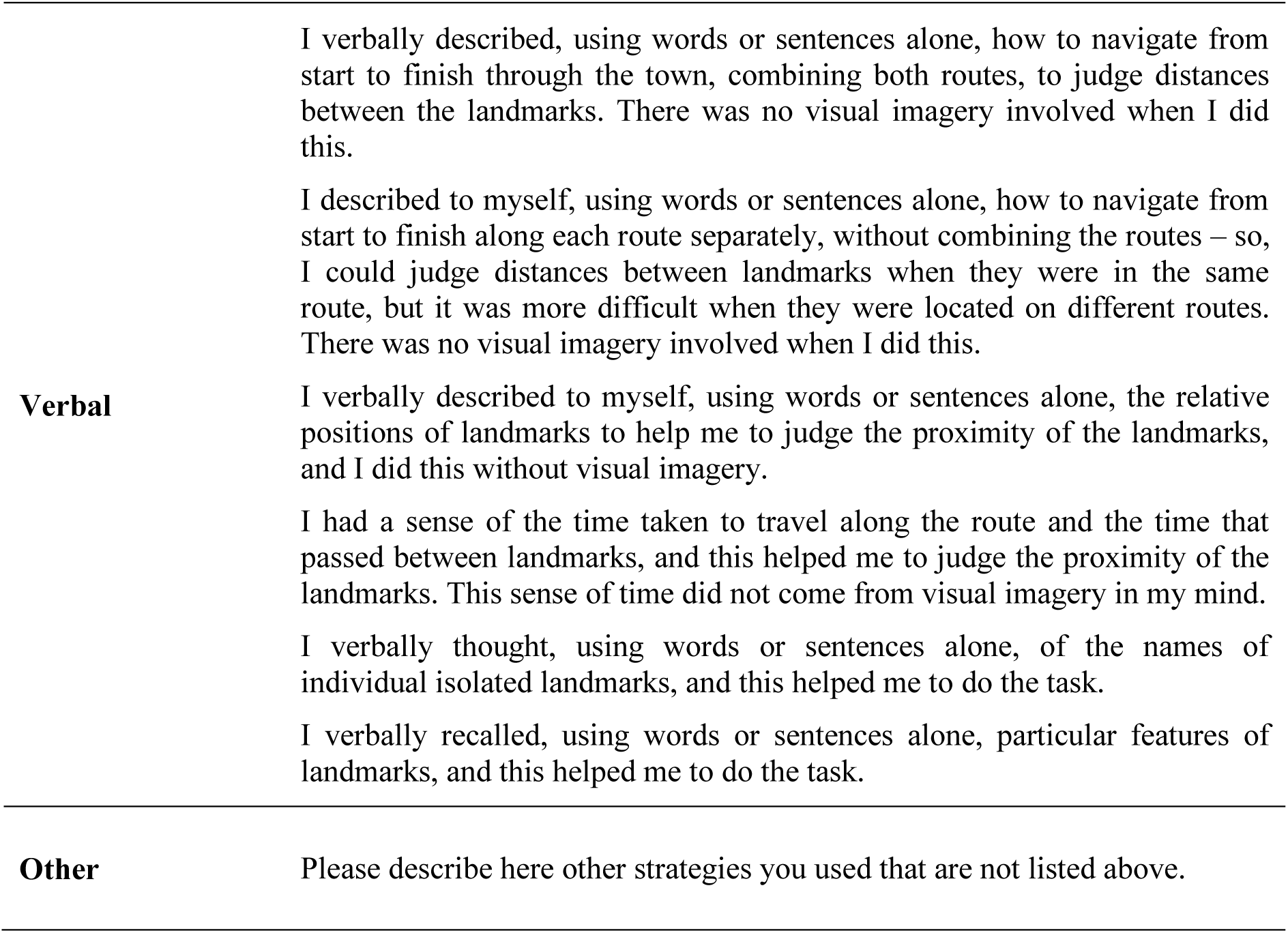

### Navigation route knowledge

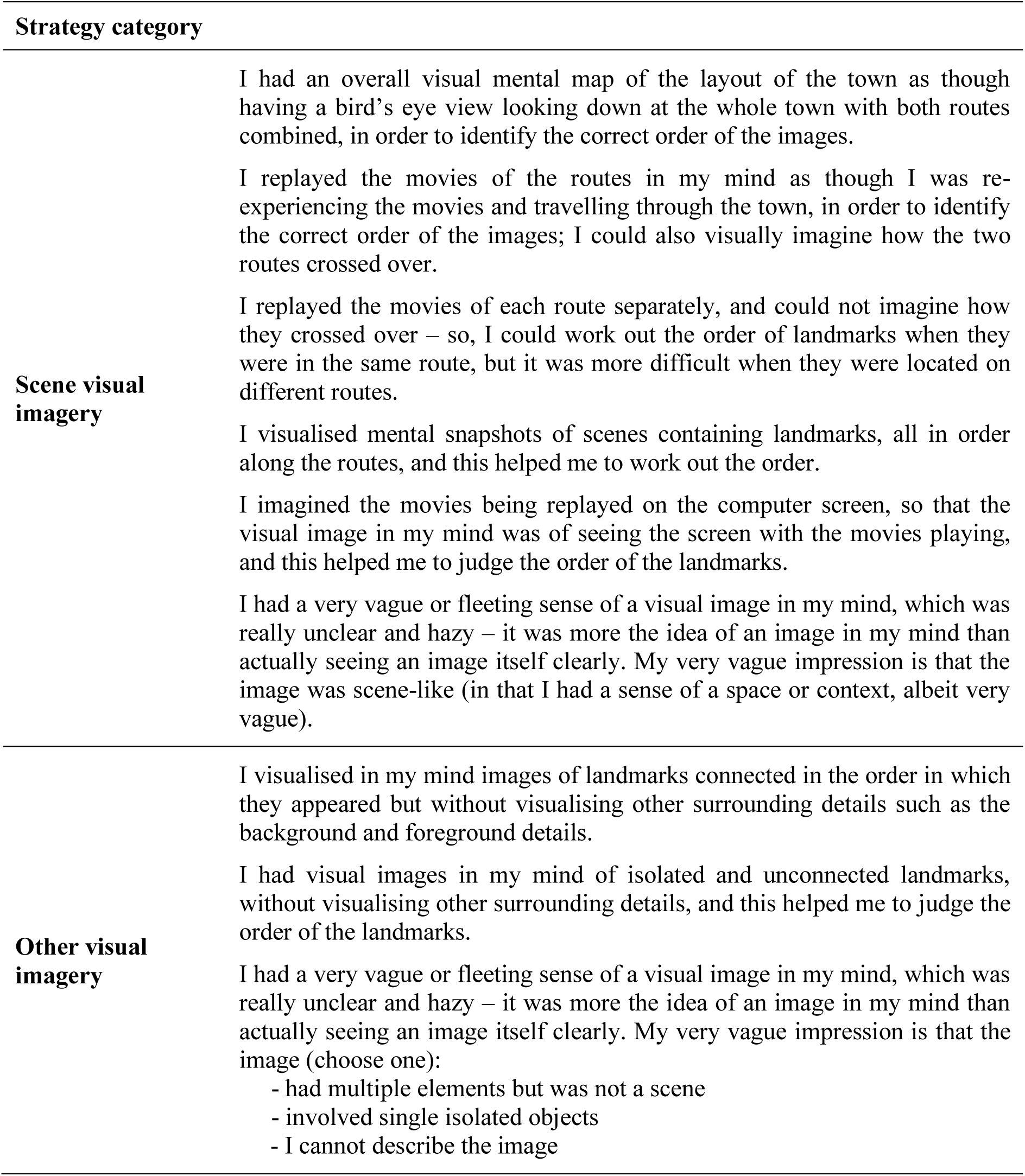

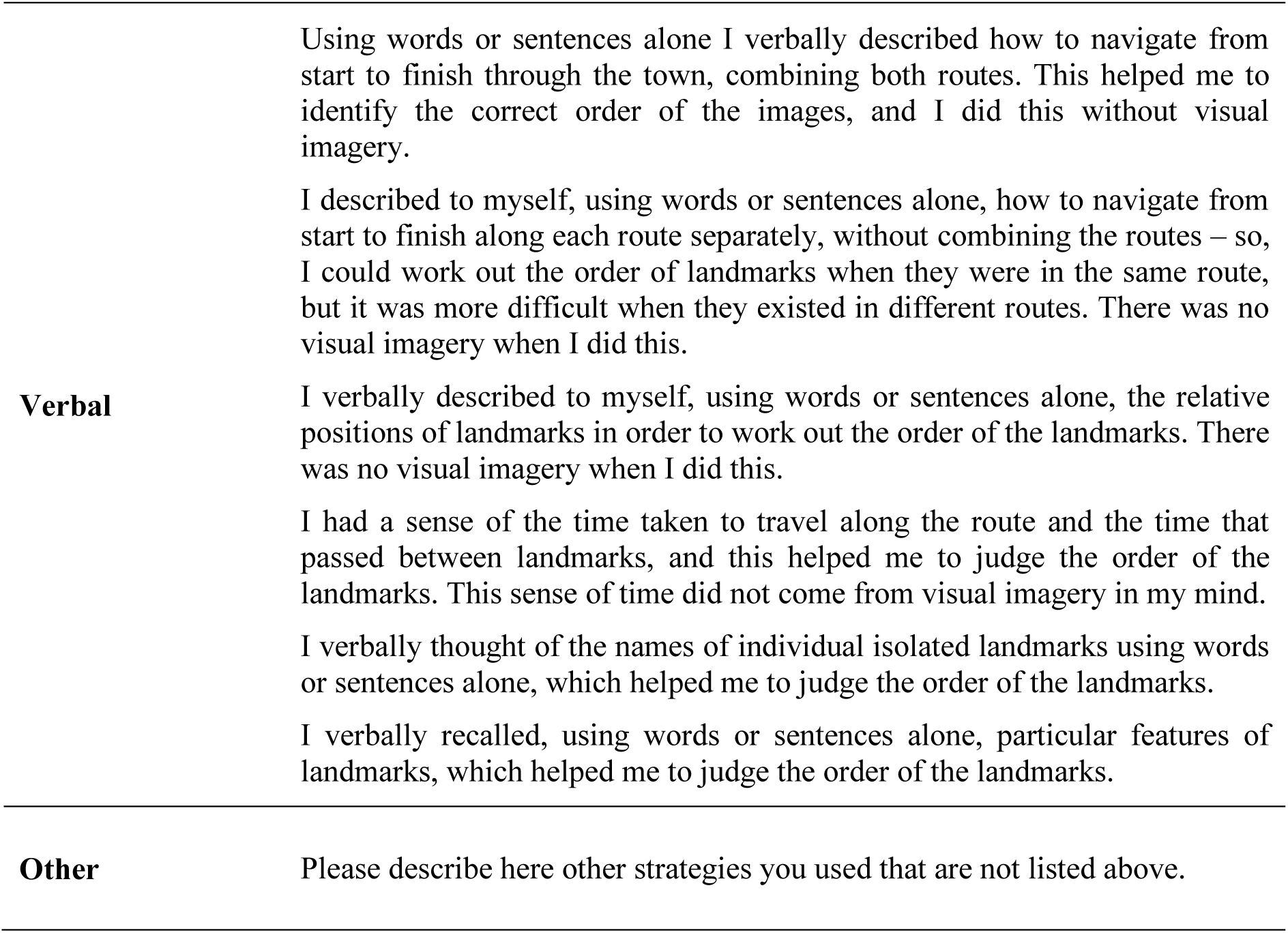

### Navigation sketch map

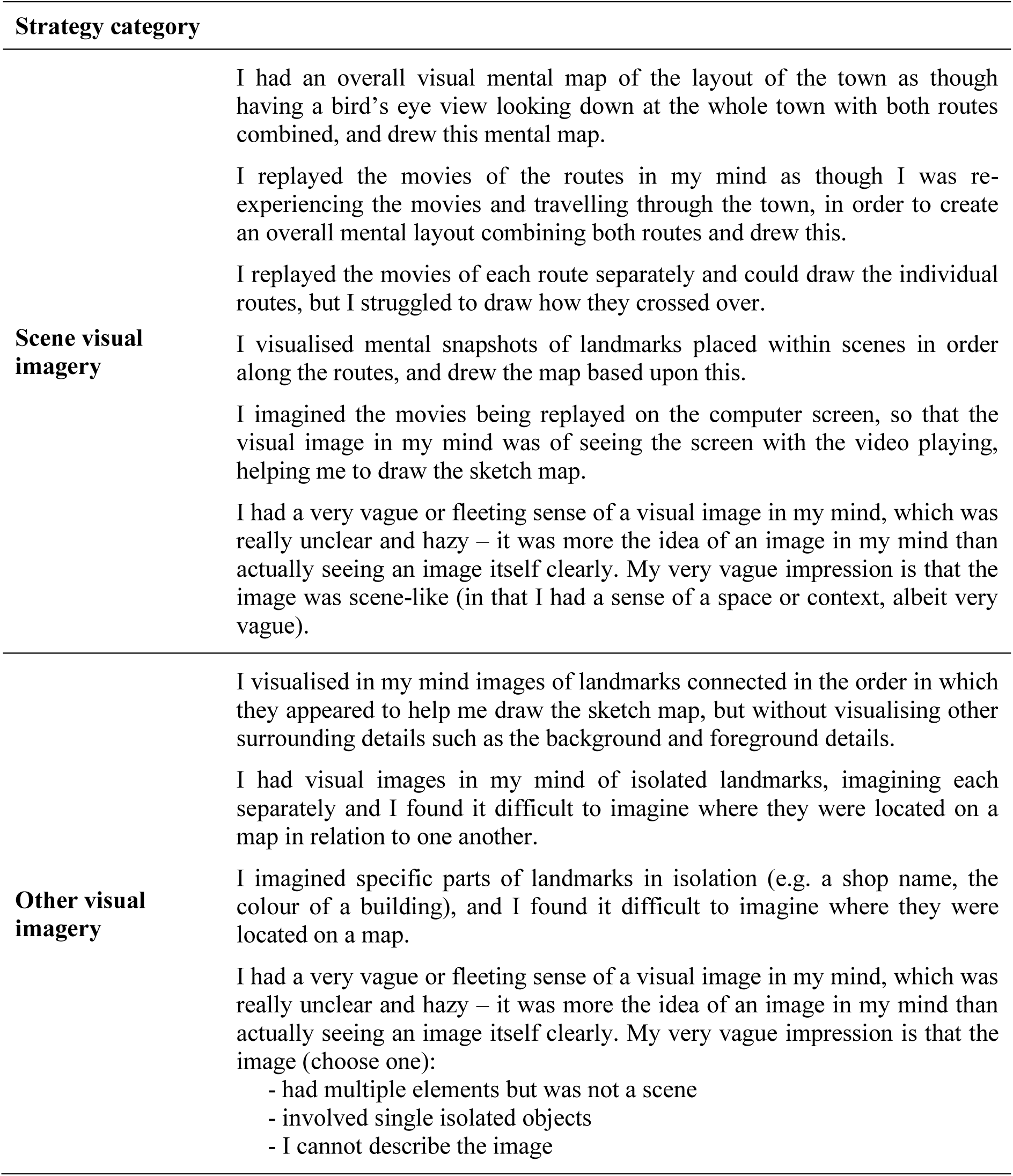

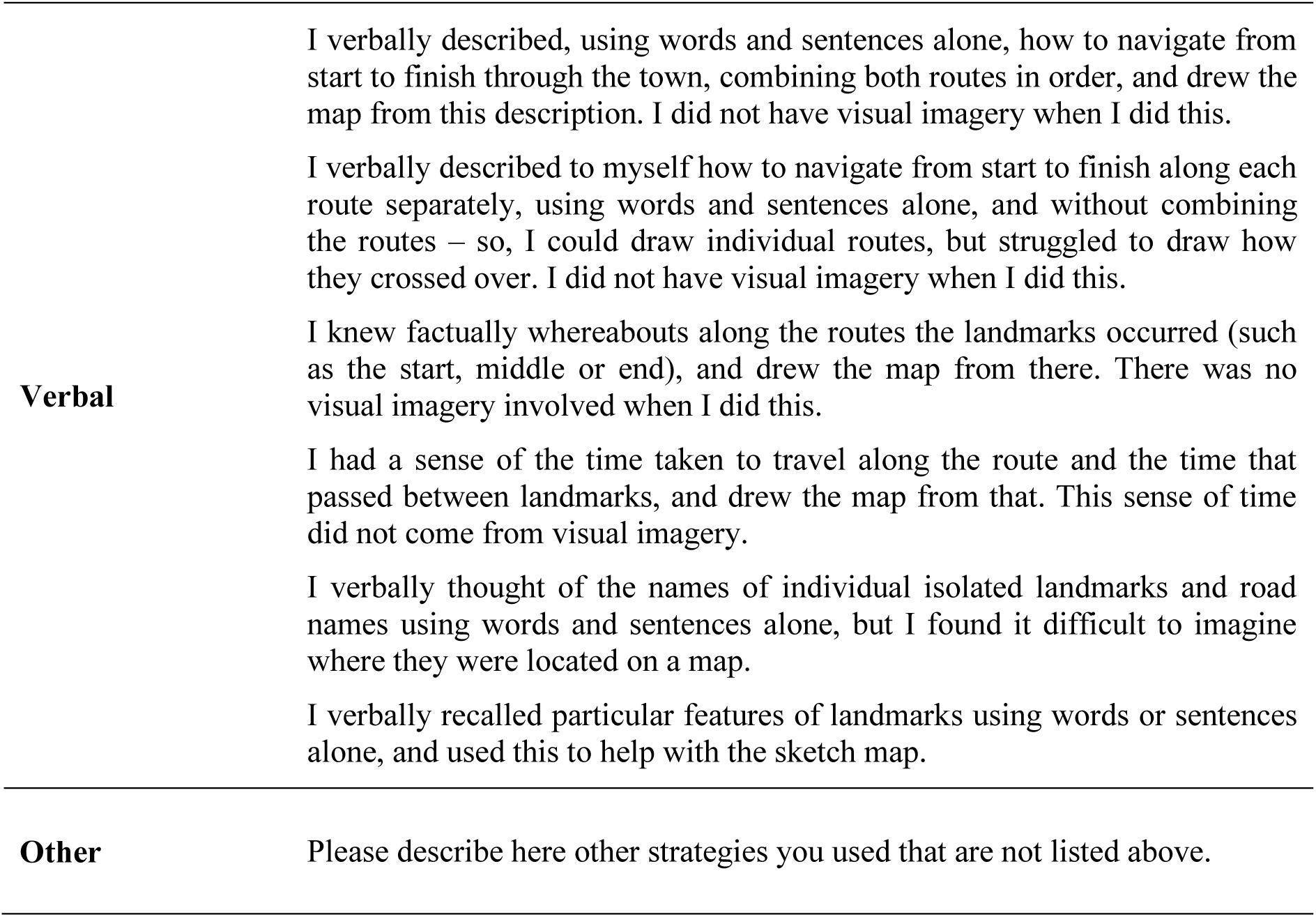

### Concrete verbal paired associates learning

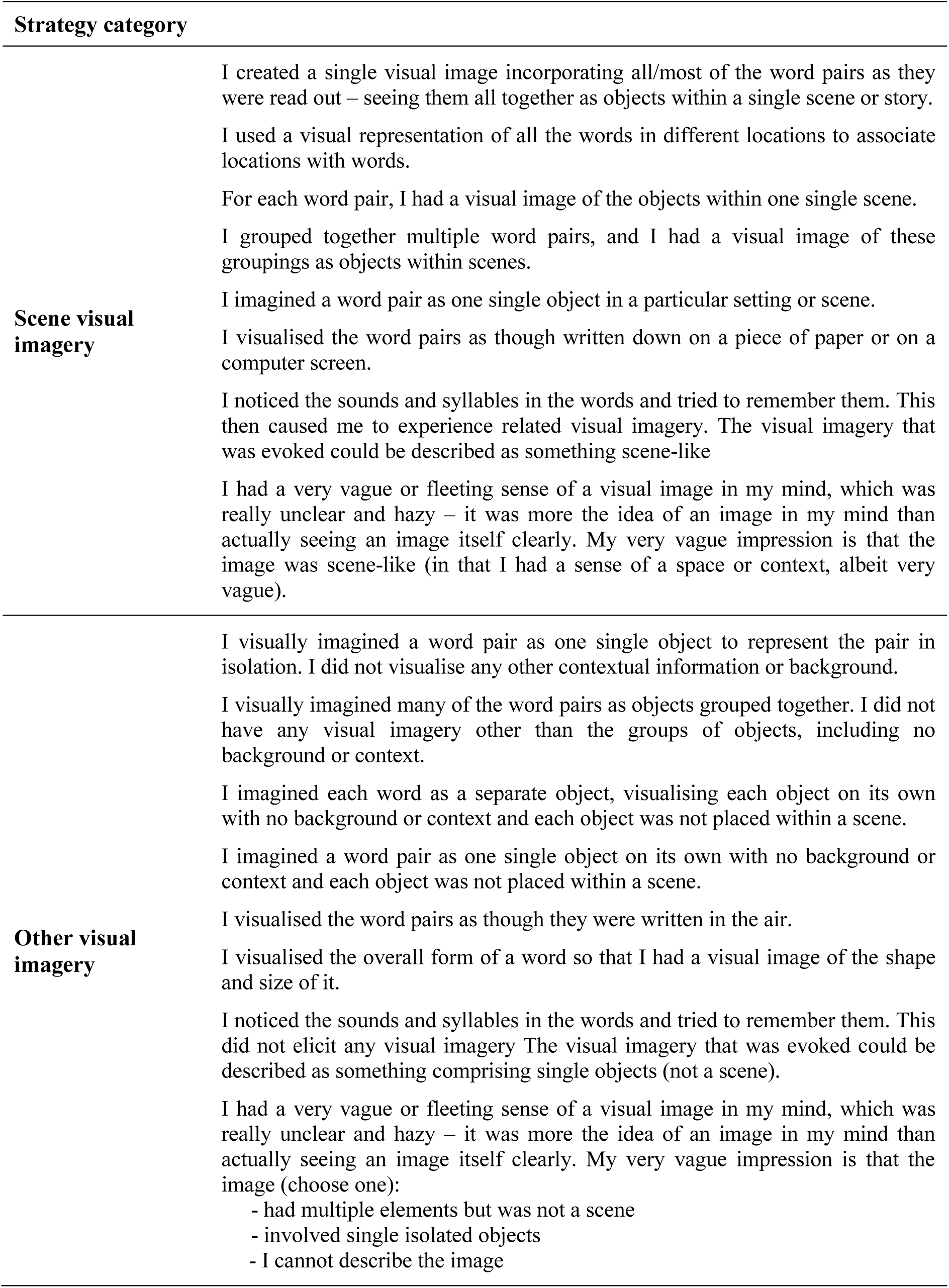

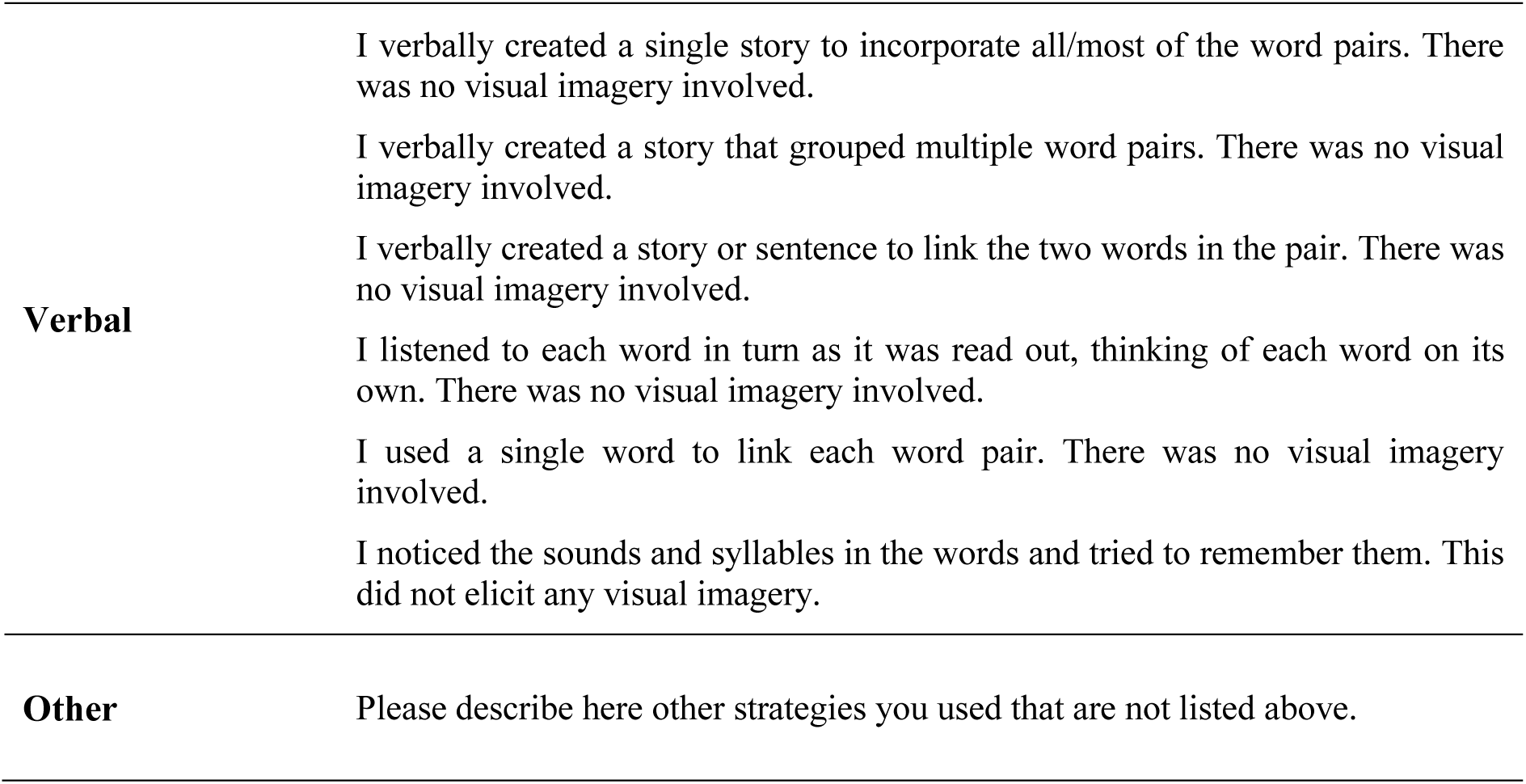

| Strategy category |  |
| --- | --- |
| **Scene visual imagery** | I created a single visual image incorporating all/most of the word pairs as they were read out – seeing them all together as objects within a single scene or story.<br>I used a visual representation of all the words in different locations to associate locations with words.<br>For each word pair, I had a visual image of the objects within one single scene.<br>I grouped together multiple word pairs, and I had a visual image of these groupings as objects within scenes.<br>I imagined a word pair as one single object in a particular setting or scene.<br>I visualised the word pairs as though written down on a piece of paper or on a computer screen.<br>I noticed the sounds and syllables in the words and tried to remember them. This then caused me to experience related visual imagery. The visual imagery that was evoked could be described as something scene-like<br>I had a very vague or fleeting sense of a visual image in my mind, which was really unclear and hazy – it was more the idea of an image in my mind than actually seeing an image itself clearly. My very vague impression is that the image was scene-like (in that I had a sense of a space or context, albeit very vague). |
| **Other visual imagery** | I visually imagined a word pair as one single object to represent the pair in isolation. I did not visualise any other contextual information or background.<br>I visually imagined many of the word pairs as objects grouped together. I did not have any visual imagery other than the groups of objects, including no background or context.<br>I imagined each word as a separate object, visualising each object on its own with no background or context and each object was not placed within a scene.<br>I imagined a word pair as one single object on its own with no background or context and each object was not placed within a scene.<br>I visualised the word pairs as though they were written in the air.<br>I visualised the overall form of a word so that I had a visual image of the shape and size of it.<br>I noticed the sounds and syllables in the words and tried to remember them. This did not elicit any visual imagery. The visual imagery that was evoked could be described as something comprising single objects (not a scene).<br>I had a very vague or fleeting sense of a visual image in my mind, which was really unclear and hazy – it was more the idea of an image in my mind than actually seeing an image itself clearly. My very vague impression is that the image (choose one):<br>- had multiple elements but was not a scene<br>- involved single isolated objects<br>- I cannot describe the image |
| **Verbal** | I verbally created a single story to incorporate all/most of the word pairs. There was no visual imagery involved.<br>I verbally created a story that grouped multiple word pairs. There was no visual imagery involved.<br>I verbally created a story or sentence to link the two words in the pair. There was no visual imagery involved.<br>I listened to each word in turn as it was read out, thinking of each word on its own. There was no visual imagery involved.<br>I used a single word to link each word pair. There was no visual imagery involved.<br>I noticed the sounds and syllables in the words and tried to remember them. This did not elicit any visual imagery. |
| **Other** | Please describe here other strategies you used that are not listed above. |

### Concrete verbal paired associates delayed recall

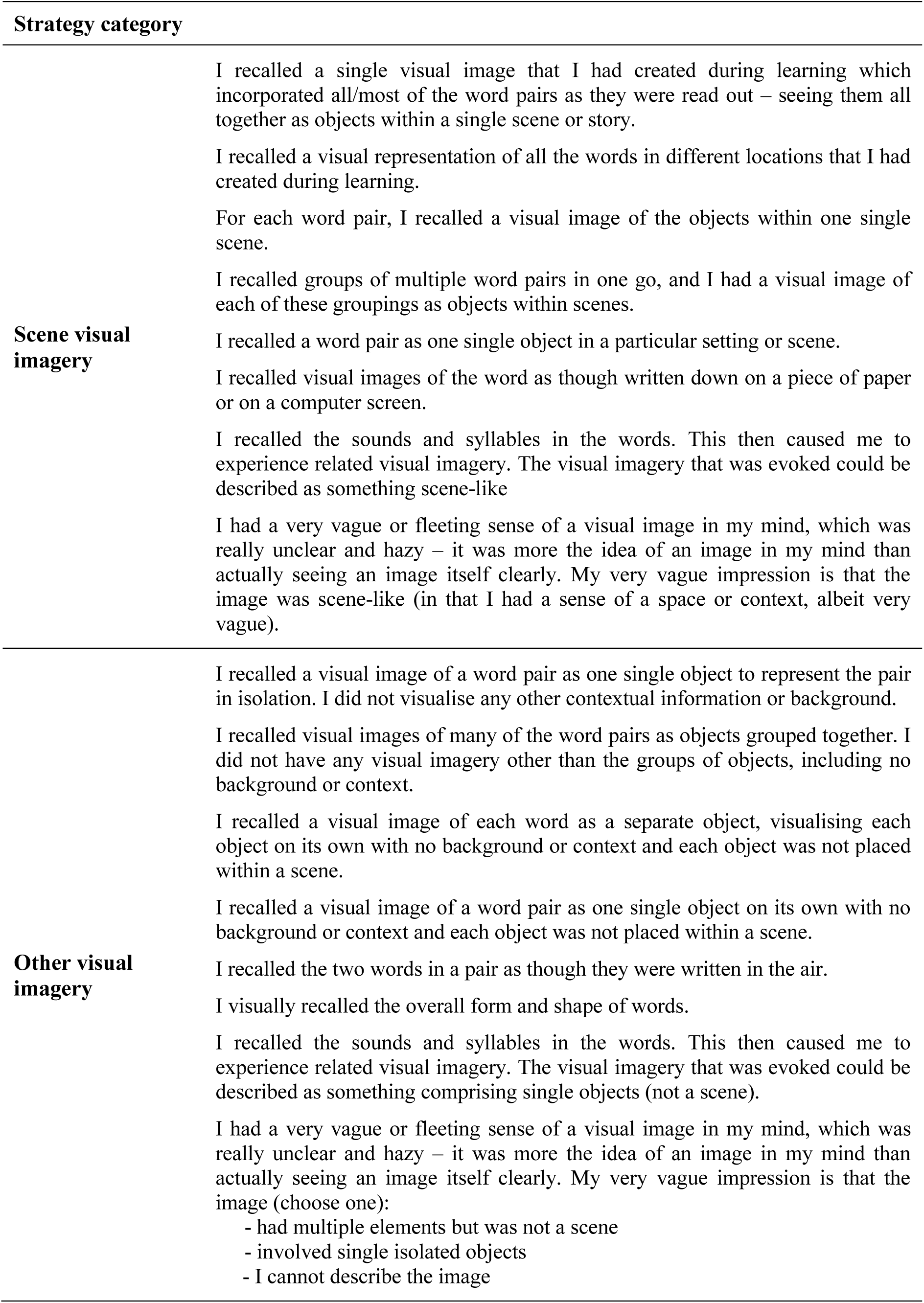

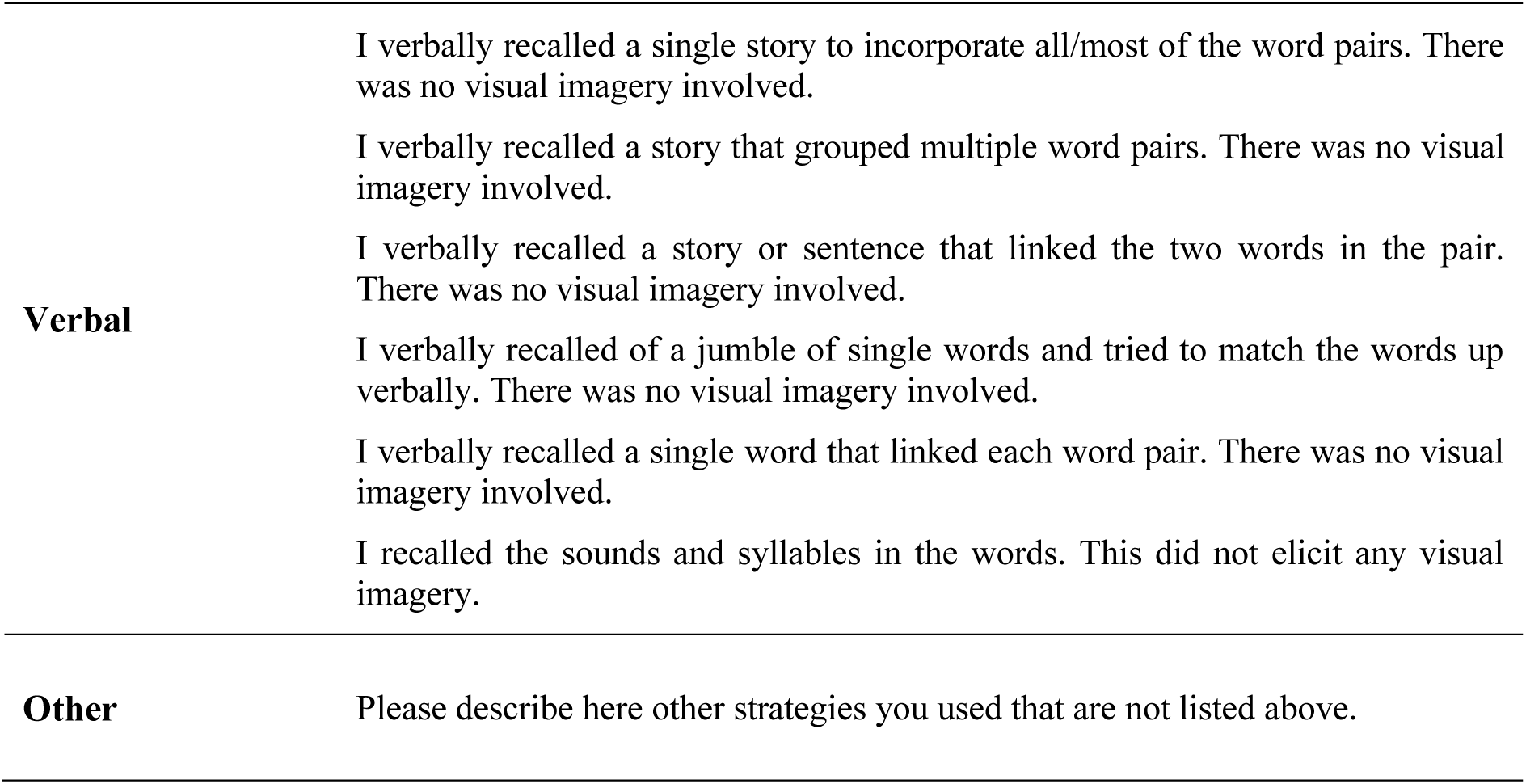

### Abstract verbal paired associates learning*

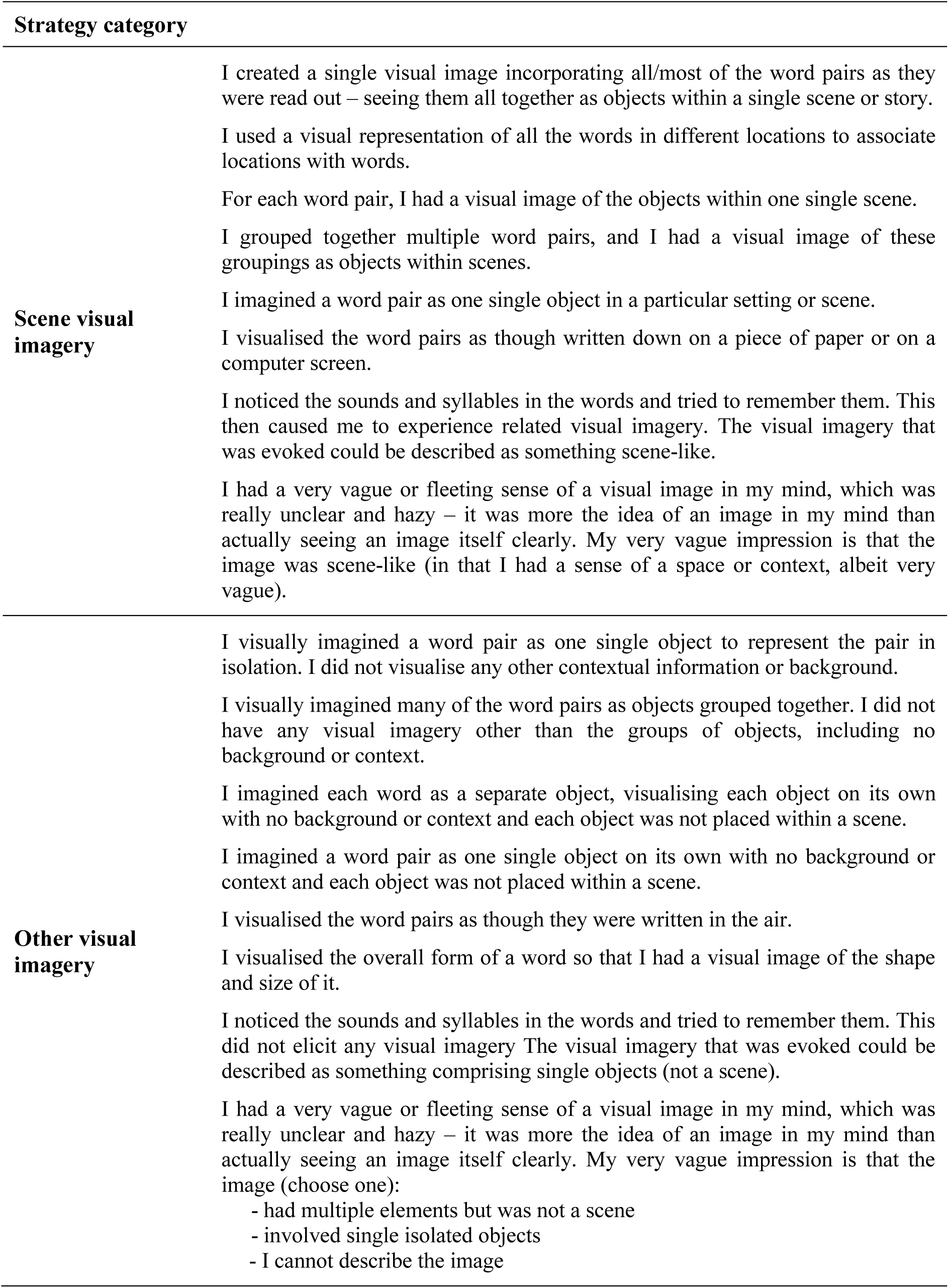

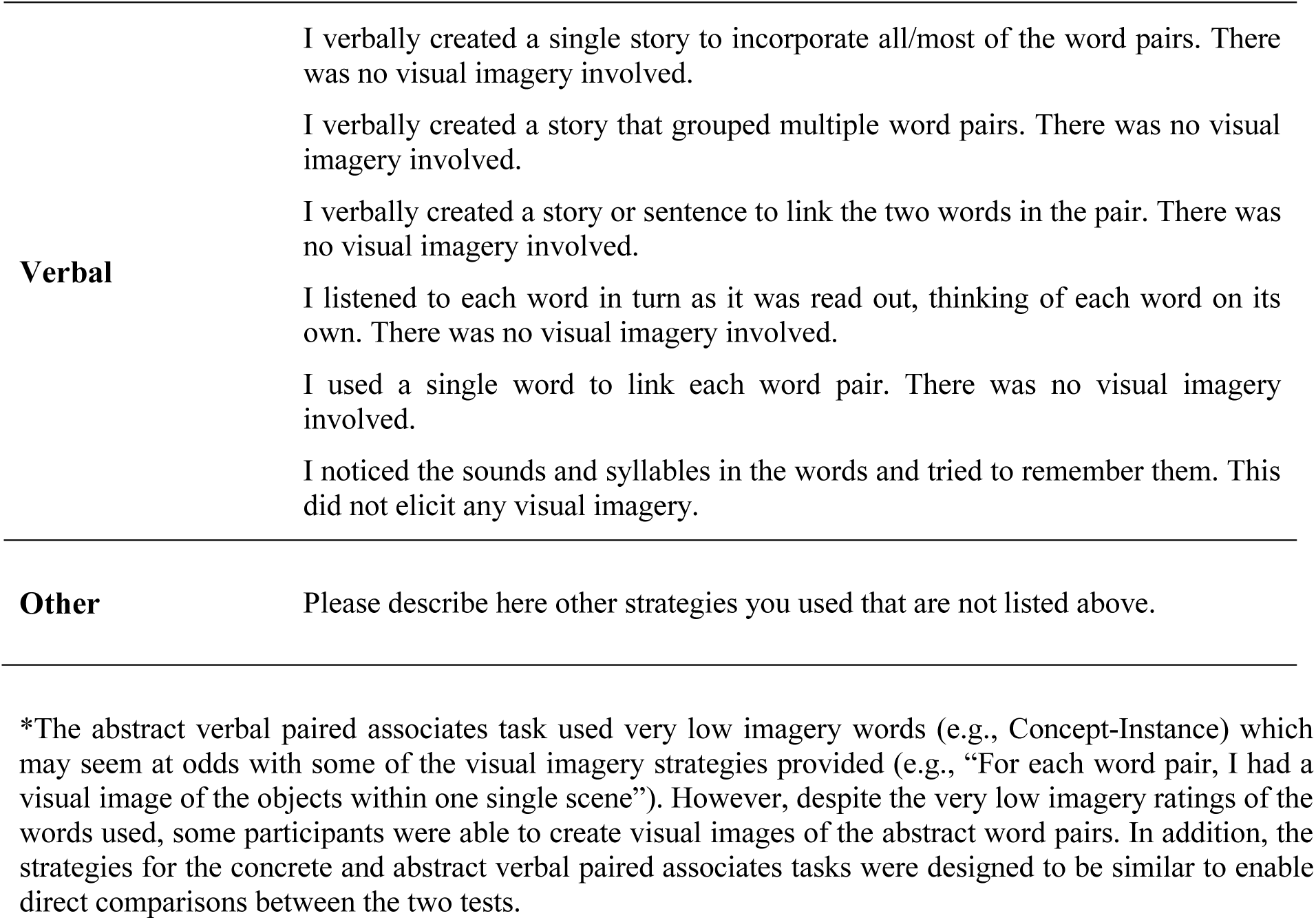

### Abstract verbal paired associates delayed recall*

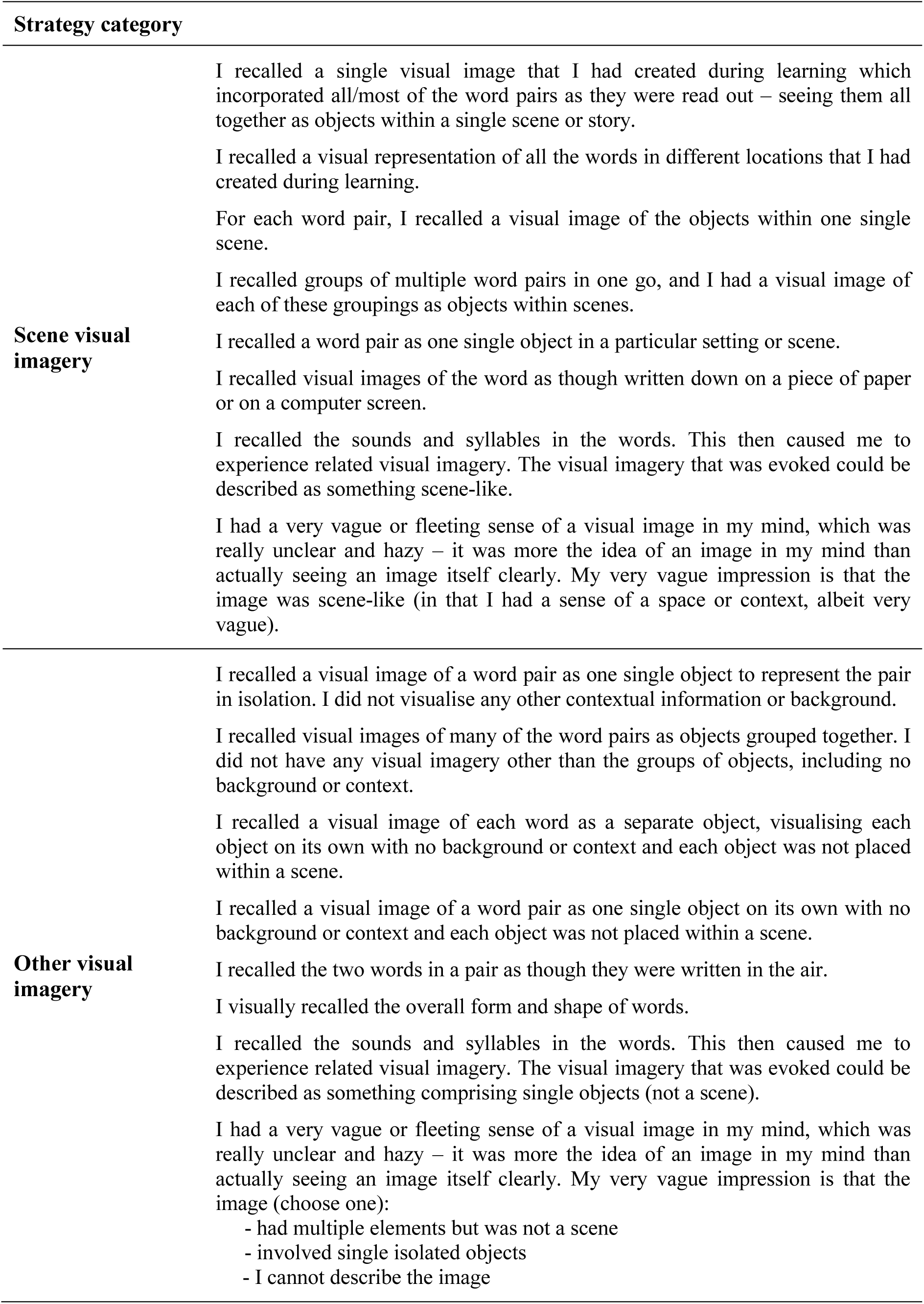

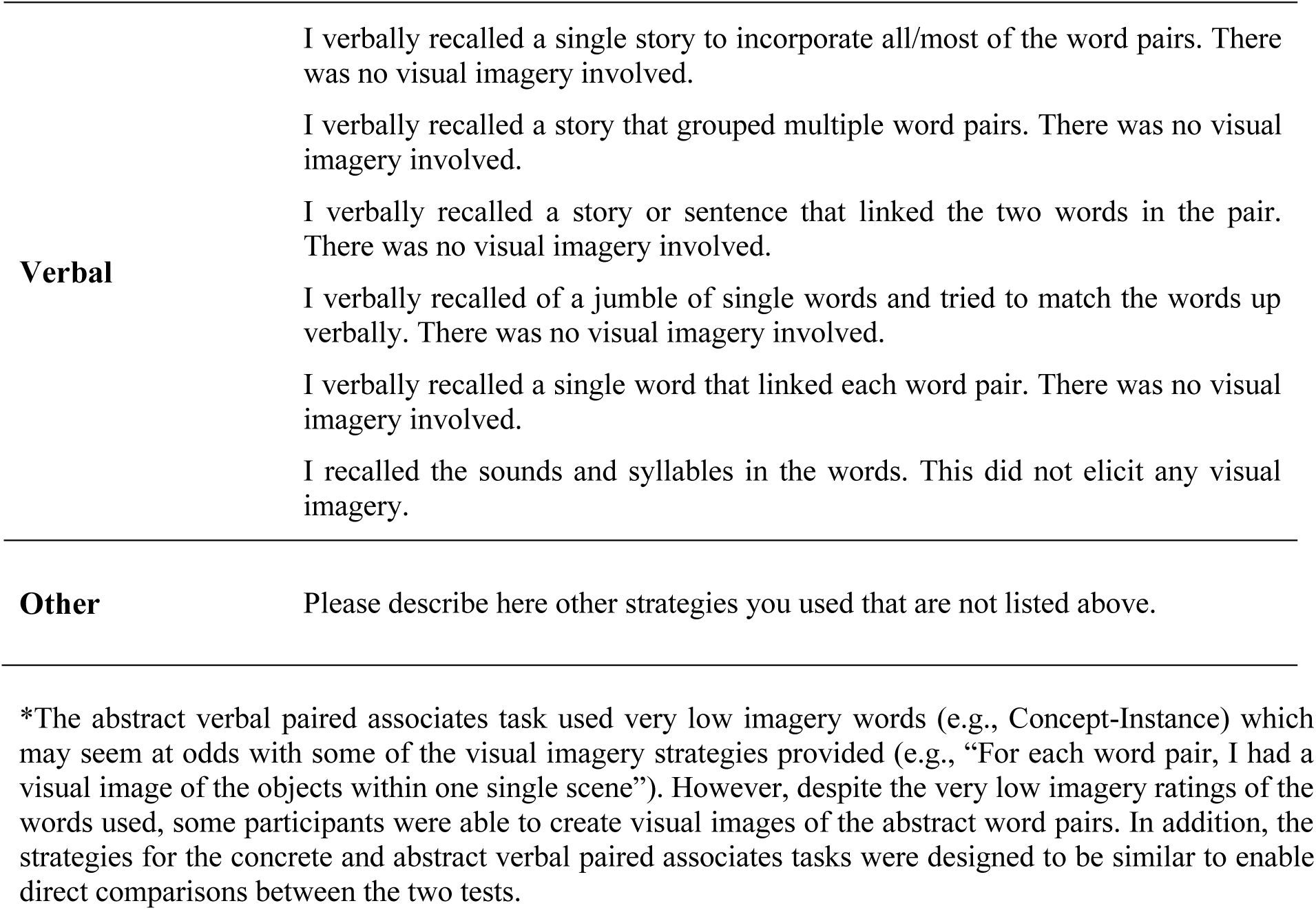

### Dead or alive task

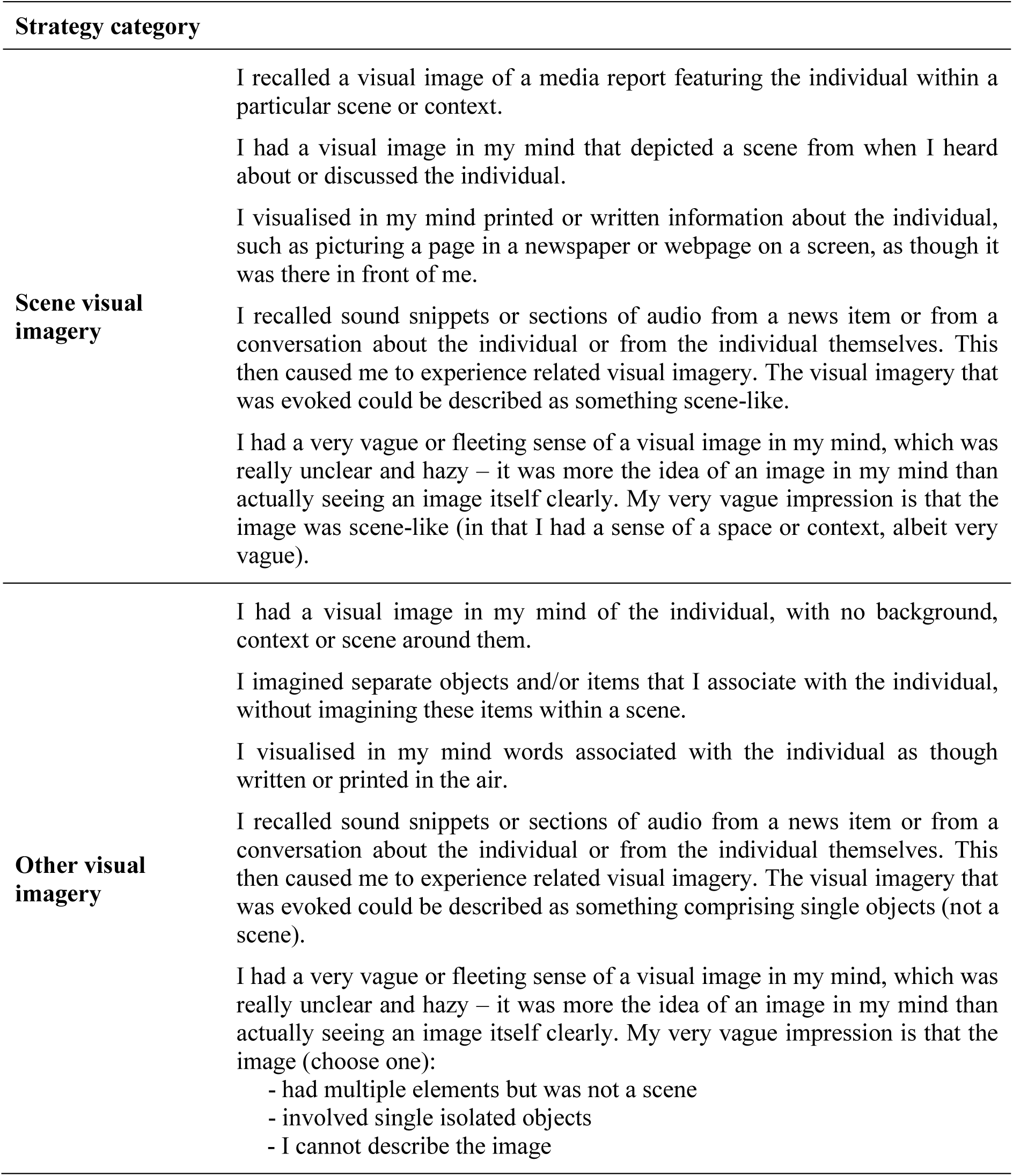

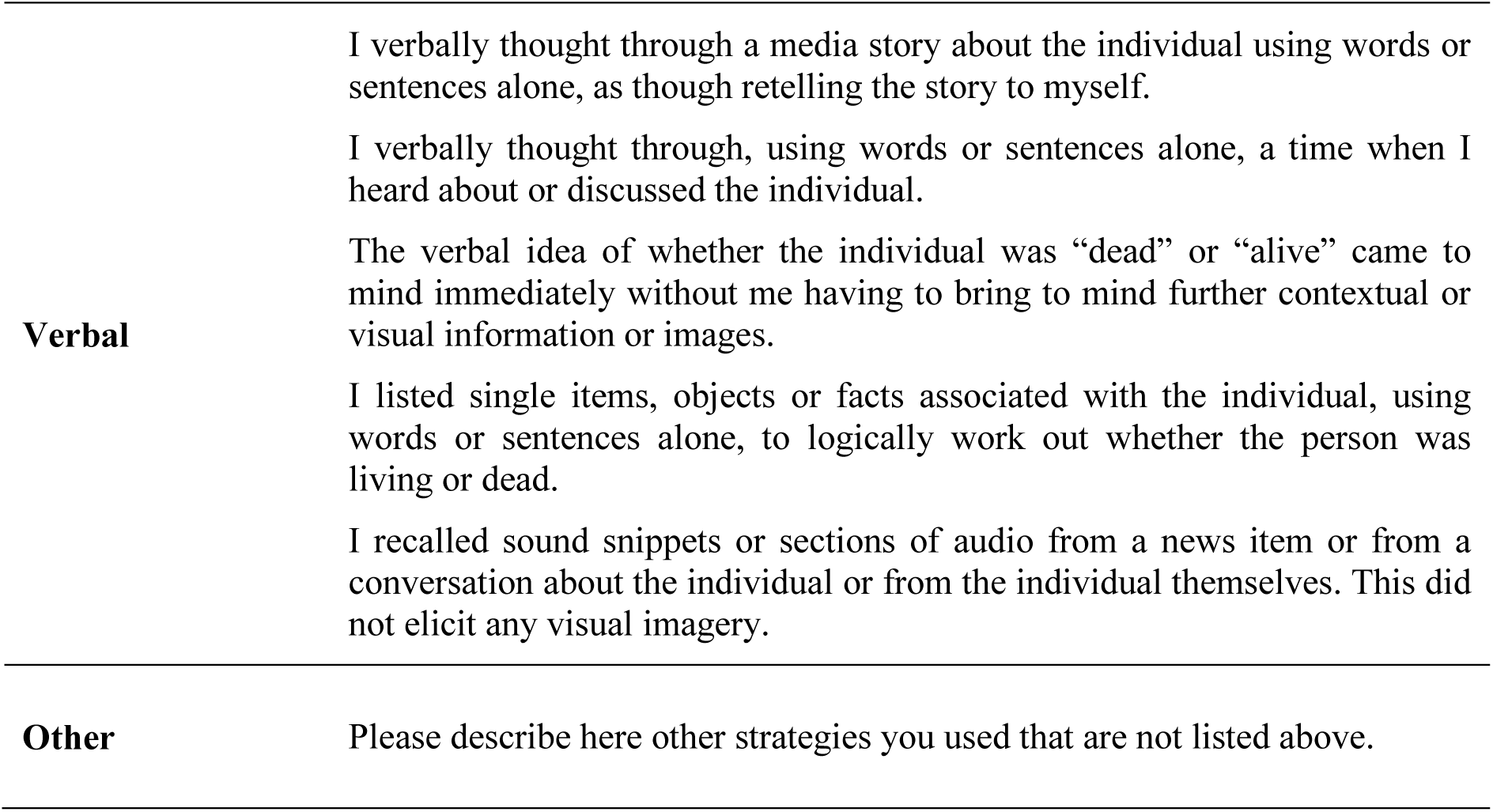

### “Other” responses

Other responses were examined to ascertain whether or not the descriptions closely resembled strategies that were already listed and, if this was the case, a strategy was reallocated from Other to the relevant strategy. Here, we detail how many rank 1 Other responses were given for each task, and to which strategy category they were reallocated if that was appropriate (scene visual imagery, other visual imagery, verbal), or whether the Other description contained no additional information that could be used for strategy classification (e.g., “I guessed”; “I imagined how I would feel in that scene”). There was no situation where the Other description referred to a new strategy that was not already represented on the list.

For the **scene construction task**, seven participants provided rank 1 Other responses. Two of these responses contained no additional information that could be used for strategy classification and five were reallocated to scene visual imagery strategies. However, all five participants whose Other responses were reallocated to scene visual imagery strategies already indicated rank 1 scene visual imagery strategies from the list provided.

On the **autobiographical interview**, for childhood autobiographical memories, four participants indicated rank 1 Other responses. Three of these responses contained no additional information that could be used for strategy classification and one was reallocated to a scene visual imagery strategy. However, this participant had already indicated a rank 1 scene visual imagery strategy from the list provided. For teenage autobiographical memories, seven participants indicated rank 1 Other responses. Six of these responses contained no additional information that could be used for strategy classification and one was reallocated to a scene visual imagery strategy. Again, however, this participant had already indicated a rank 1 scene visual imagery strategy from the list provided. For adulthood autobiographical memories, six participants indicated rank 1 Other responses, none of which contained additional information that could be used for strategy classification. For autobiographical memories from the last year, five participants indicated rank 1 Other responses, none of which contained additional information that could be used for strategy classification.

For the **future thinking task**, four participants indicated rank 1 Other responses. Three of these responses contained no additional information that could be used for strategy classification and one was reallocated to a scene visual imagery strategy. However, this participant had already indicated a rank 1 scene visual imagery strategy from the list provided.

For the **navigation tasks**, when viewing the movies while learning, four participants indicated rank 1 Other responses. One of these responses contained no additional information that could be used for strategy classification, one was reallocated to a scene visual imagery strategy, one to other visual imagery strategy and one to a verbal strategy. However, all three participants whose other responses were reallocated had already indicated rank 1 strategies in the corresponding strategy category. For the navigation movie clip recognition task, six participants indicated rank 1 Other responses. Four of these responses contained no additional information that could be used for strategy classification, one was reallocated to a scene visual imagery strategy and one to other visual imagery strategy. Again, both the participants whose strategies were reallocated had already indicated rank 1 strategies in the corresponding strategy category. For the navigation scene recognition task, six participants indicated rank 1 Other responses. Four of these responses contained no additional information that could be used for strategy classification, one was reallocated to a scene visual imagery strategy and one to a verbal strategy. For the participant whose Other response was reallocated to a scene visual imagery strategy, they had already indicated a rank 1 scene visual imagery strategy. However, the participant whose Other response was reallocated to a verbal strategy had not indicated using a rank 1 verbal strategy, even though their Other strategy description did match one of the verbal strategies on the list provided. For the navigation proximity judgements task, five participants indicated rank 1 Other responses. Four of these responses contained no additional information that could be used for strategy classification and one was reallocated to other visual imagery strategy. While this participant had not indicated using a rank 1 other visual imagery strategy, the strategy detailed did match other visual imagery strategy from the strategy list provided. For the navigation route knowledge task, five participants indicated rank 1 Other responses, none of which contained additional information that could be used for strategy classification. For the navigation sketch map, three participants indicated rank 1 Other responses, none of which contained additional information that could be used for strategy classification.

For the **concrete verbal paired associates** (VPA) learning task, seven participants indicated rank 1 Other responses. Three of these responses contained no additional information that could be used for strategy classification and four were reallocated to verbal strategies. However, all four participants whose Other responses were reallocated to verbal strategies already indicated rank 1verbal strategies from the list provided. For the concrete VPA delayed recall task, eight participants indicated rank 1 Other responses. Six of the responses contained no additional information that could be used for strategy classification and two were reallocated to verbal strategies. Again, both participants whose Other responses were reallocated to verbal strategies had already indicated rank 1 verbal strategies from the list provided.

For the **abstract VPA** learning task, 15 participants indicated rank 1 Other responses. Four of these responses contained no additional information that could be used for strategy classification, two were reallocated to scene visual imagery strategies, four to other visual imagery strategies and five to verbal strategies. However, all the participants whose strategies were reallocated had already indicated rank 1 strategies in the corresponding strategy category. For the abstract VPA delayed recall task, 15 participants indicated rank 1 Other responses. Ten of these responses contained no additional information that could be used for strategy classification, two were reallocated to scene visual imagery strategies, two to other visual imagery strategies and one to a verbal strategy. As before, all the participants whose strategies were reallocated had already indicated rank 1 strategies in the corresponding strategy category.

Finally, for the **dead or alive** semantic memory task, two participants indicated rank 1 Other responses. One of these responses contained no additional information that could be used for strategy classification while the other was reallocated to other visual imagery strategy. However, this participant had already indicated using a rank 1 other visual imagery strategy from the list provided.

Overall, therefore, examination of the Other responses resulted in very little additional information not already identified by the strategy questionnaires.

## REFERENCES

Addis, D. R., Wong, A. T. and Schacter, D. L. (2007). Remembering the past and imagining the future: Common and distinct neural substrates during event construction and elaboration. Neuropsychologia, 45(7), 1363–1377. doi: http://dx.doi.org/10.1016/j.neuropsychologia.2006.10.016

Andelman, F., Hoofien, D., Goldberg, I., Aizenstein, O. and Neufeld, M. Y. (2010). Bilateral hippocampal lesion and a selective impairment of the ability for mental time travel. Neurocase, 16(5), 426–435. doi: http://dx.doi.org/10.1080/13554791003623318

Andrews-Hanna, J. R., Reidler, J. S., Sepulcre, J., Poulin, R. and Buckner, R. L. (2010). Functional-anatomic fractionation of the brain’s default network. Neuron, 65(4), 550–562. doi: http://doi.org/10.1016/j.neuron.2010.02.005

Andrews-Hanna, J. R., Saxe, R. and Yarkoni, T. (2014). Contributions of episodic retrieval and mentalizing to autobiographical thought: Evidence from functional neuroimaging, resting-state connectivity, and fMRI meta-analyses. Neuroimage, 91, 324–335. doi: http://dx.doi.org/10.1016/j.neuroimage.2014.01.032

Barry, D. N., Barnes, G. R., Clark, I. A. and Maguire, E. A. (2018). The neural dynamics of novel scene imagery. J. Neurosci., 32(22), 4375–4386. doi: http://dx.doi.org/10.1523/JNEUROSCI.2497-18.2019

Binder, J. R. and Desai, R. H. (2011). The neurobiology of semantic memory. Trends Cogn. Sci., 15(11), 527–536. doi: http://dx.doi.org/10.1016/j.tics.2011.10.001

Boltwood, C. E. and Blick, K. A. (1970). The delineation and application of three mnemonic techniques. Psychon. Sci., 20(6), 339–341. doi: http://dx.doi.org/10.3758/BF03335678

Bower, G. (1970). Imagery as a relational organiser in in associative learning. J Verbal Learning Verbal Behav, 9, 529–533. doi: http://dx.doi.org/10.1016/s0022-5371(70)80096-2

Clark, I. A., Hotchin, V., Monk, A., Pizzamiglio, G., Liefgreen, A. and Maguire, E. A. (2019). Identifying the cognitive processes underpinning hippocampal-dependent tasks. J. Exp. Psychol. Gen., 148(11), 1861–1881. doi: http://dx.doi.org/10.1037/xge0000582

Clark, I. A., Kim, M. and Maguire, E. A. (2018). Verbal paired associates and the hippocampus: The role of scenes. J. Cogn. Neurosci., 30(12), 1821–1845. doi: http://dx.doi.org/10.1162/jocn_a_01315

Clark, I. A. and Maguire, E. A. (2016). Remembering preservation in hippocampal amnesia. Annu. Rev. Psychol., 67(1), 51–82. doi: http://dx.doi.org/10.1146/annurev-psych-122414-033739

Clark, I. A. and Maguire, E. A. (2020). Do questionnaires reflect their purported cognitive functions? Cognition, 195, 104114. doi: http://dx.doi.org/10.1016/j.cognition.2019.104114

Dalton, M. A., Zeidman, P., McCormick, C. and Maguire, E. A. (2018). Differentiable processing of objects, associations and scenes within the hippocampus. J. Neurosci., 38(38), 8146–8159. doi: http://dx.doi.org/10.1523/JNEUROSCI.0263-18.2018

Dunlosky, J. and Hertzog, C. (1998). Aging and deficits in associative memory: What is the role of strategy production? Psychol. Aging, 13(4), 597–607. doi: http://dx.doi.org/10.1037/0882-7974.13.4.597

Dunlosky, J. and Hertzog, C. (2001). Measuring strategy production during associative learning: The relative utility of concurrent versus retrospective reports. Mem. Cognit., 29(2), 247–253. doi: http://dx.doi.org/10.3758/BF03194918

Dunlosky, J. and Kane, M. J. (2007). The contributions of strategy use to working memory span: A comparison of strategy assessment methods. Q. J. Exp. Psychol., 60(9), 1227–1245. doi: http://dx.doi.org/10.1080/17470210600926075

Einstein, G. O. and McDaniel, M. A. (1987). Distinctiveness and the mnemonic benefits of bizarre imagery. In M. A. McDaniel & M. Pressley (Eds.), Imagery and related mnemonic processes (pp. 78–102). New York: Springer. doi: http://dx.doi.org/10.1007/978-1-4612-4676-3_4

Ekstrom, A. D., Kahana, M. J., Caplan, J. B., Fields, T. A., Isham, E. A., Newman, E. L., et al. (2003). Cellular networks underlying human spatial navigation. Nature, 425(6954), 184–188. doi: http://www.nature.com/nature/journal/v425/n6954/suppinfo/nature01964_S1.html

Ekstrom, A. D. and Ranganath, C. (2018). Space, time, and episodic memory: The hippocampus is all over the cognitive map. Hippocampus, 28(9), 680–687. doi: http://dx.doi.org/10.1002/hipo.22750

Ericsson, K. A. and Simon, H. A. (1980). Verbal reports as data. Psychol. Rev., 87(3), 215–251. doi: http://dx.doi.org/10.1037/0033-295X.87.3.215

Giovanello, K. S., Verfaellie, M. and Keane, M. M. (2003). Disproportionate deficit in associative recognition relative to item recognition in global amnesia. Cogn. Affect. Behav. Neurosci., 3(3), 186–194. doi: http://dx.doi.org/10.3758/cabn.3.3.186

Greenberg, D. L. and Knowlton, B. J. (2014). The role of visual imagery in autobiographical memory. Mem. Cognit., 42(6), 922–934. doi: http://dx.doi.org/10.3758/s13421-014-0402-5

Hassabis, D. and Maguire, E. A. (2007). Deconstructing episodic memory with construction. Trends Cogn. Sci., 11(7), 299–306. doi: http://dx.doi.org/10.1016/j.tics.2007.05.001

Hassabis, D., Kumaran, D., Vann, S. D. and Maguire, E. A. (2007a). Patients with hippocampal amnesia cannot imagine new experiences. Proc. Natl. Acad. Sci. USA, 104(5), 1726–1731. doi: http://dx.doi.org/10.1073/pnas.0610561104

Hassabis, D., Kumaran, D. and Maguire, E. A. (2007b). Using imagination to understand the neural basis of episodic memory. J. Neurosci., 27(52), 14365–14374. doi: http://dx.doi.org/10.1523/jneurosci.4549-07.2007

Hertzog, C., McGuire, C. L. and Lineweaver, T. T. (1998). Aging, attributions, perceived control, and strategy use in a free recall task. Aging Neuropsychol. Cogn., 5(2), 85–106. doi: http://dx.doi.org/10.1076/anec.5.2.85.601

Kapur, N., Young, A., Bateman, D. and Kennedy, P. (1989). Focal retrograde amnesia: A long term clinical and neuropsychological follow-up. Cortex, 25(3), 387–402. doi: http://dx.doi.org/10.1016/S0010-9452(89)80053-X

Klein, S. B., Loftus, J. and Kihlstrom, J. F. (2002). Memory and temporal experience: The effects of episodic memory loss on an amnesic patient’s ability to remember the past and imagine the future. Soc. Cognition, 20(5), 353–379. doi: http://dx.doi.org/10.1521/soco.20.5.353.21125

Kroll, N. E. A., Schepeler, E. M. and Angin, K. T. (1986). Bizarre imagery: The misremembered mnemonic. J. Exp. Psychol. Learn. Mem. Cogn., 12(1), 42–53. doi: http://dx.doi.org/10.1037//0278-7393.12.1.42

Levine, B., Svoboda, E., Hay, J. F., Winocur, G. and Moscovitch, M. (2002). Aging and autobiographical memory: Dissociating episodic from semantic retrieval. Psychol. Aging, 17(4), 677–689. doi: http://dx.doi.org/10.1037/0882-7974.17.4.677

Logie, R. H., Sala, S. D., Laiacona, M., Chalmers, P. and Wynn, V. (1996). Group aggregates and individual reliability: The case of verbal short-term memory. Mem. Cognit., 24(3), 305–321. doi: http://dx.doi.org/10.3758/BF03213295

Maguire, E. A., Gadian, D. G., Johnsrude, I. S., Good, C. D., Ashburner, J., Frackowiak, R. S. J., et al. (2000). Navigation-related structural change in the hippocampi of taxi drivers. Proc. Natl. Acad. Sci. USA, 97(8), 4398–4403. doi: http://dx.doi.org/10.1073/pnas.070039597

Maguire, E. A. and Mullally, S. L. (2013). The hippocampus: A manifesto for change. J. Exp. Psychol. Gen., 142(4), 1180–1189. doi: http://dx.doi.org/10.1037/a0033650

Marschark, M. and Hunt, R. R. (1989). A reexamination of the role of imagery in learning and memory. J. Exp. Psychol. Learn. Mem. Cogn., 15(4), 710–720. doi: http://dx.doi.org/10.1037//0278-7393.15.4.710

McDaniel, M. A. and Einstein, G. O. (1986). Bizarre imagery as an effective memory aid: The importance of distinctiveness. J. Exp. Psychol. Learn. Mem. Cogn., 12(1), 54–65. doi: http://dx.doi.org/10.1037//0278-7393.12.1.54

McDaniel, M. A. and Kearney, E. M. (1984). Optimal learning strategies and their spontaneous use: The importance of task-appropriate processing. Mem. Cognit., 12(4), 361–373. doi: http://dx.doi.org/10.3758/BF03198296

Moscovitch, M., Cabeza, R., Winocur, G. and Nadel, L. (2016). Episodic memory and beyond: The hippocampus and neocortex in transformation. Annu. Rev. Psychol., 67(1), 105–134. doi: http://dx.doi.org/doi:10.1146/annurev-psych-113011-143733

Moser, E. I., Kropff, E. and Moser, M.-B. (2008). Place cells, grid cells, and the brain’s spatial representation system. Annu. Rev. Neurosci., 31(1), 69–89. doi: http://dx.doi.org/10.1146/annurev.neuro.31.061307.090723

Mullally, S. L., Intraub, H. and Maguire, E. A. (2012). Attenuated boundary extension produces a paradoxical memory advantage in amnesic patients. Curr. Biol., 22(4), 261–268. doi: http://dx.doi.org/10.1016/j.cub.2012.01.001

O’Keefe, J. and Nadel, L. (1978). The hippocampus as a cognitive map. Oxford: Clarendon Press.

Paivio, A. (1969). Mental imagery in associative learning and memory. Psychol. Rev., 76(3), 241–263. doi: http://dx.doi.org/10.1037/h0027272

Palombo, D. J., Hayes, S. M., Peterson, K. M., Keane, M. M. and Verfaellie, M. (2018). Medial temporal lobe contributions to episodic future thinking: Scene construction or future projection? Cereb. Cortex, 28(2), 447–458. doi: http://dx.doi.org/10.1093/cercor/bhw381

Race, E., Keane, M. M. and Verfaellie, M. (2011). Medial temporal lobe damage causes deficits in episodic memory and episodic future thinking not attributable to deficits in narrative construction. J. Neurosci., 31(28), 10262–10269. doi: http://dx.doi.org/10.1523/jneurosci.1145-11.2011

Reisberg, D., Pearson, D. G. and Kosslyn, S. M. (2003). Intuitions and introspections about imagery: The role of imagery experience in shaping an investigator’s theoretical views. Appl. Cogn. Psychol., 17(2), 147–160. doi: http://dx.doi.org/10.1002/acp.858

Roberts, W. A. (1968). Alphabetic coding and individual differences in modes of organization in free-recall learning. Am. J. Psychol., 81(3), 433–438. doi: http://dx.doi.org/10.2307/1420642

Robin, J. (2018). Spatial scaffold effects in event memory and imagination. Wiley Interdiscip. Rev. Cogn. Sci., e1462. doi: http://dx.doi.org/10.1002/wcs.1462

Rosenbaum, R. S., Gilboa, A., Levine, B., Winocur, G. and Moscovitch, M. (2009). Amnesia as an impairment of detail generation and binding: Evidence from personal, fictional, and semantic narratives in K.C. Neuropsychologia, 47(11), 2181–2187. doi: http://dx.doi.org/10.1016/j.neuropsychologia.2008.11.028

Rubin, D. C. and Umanath, S. (2015). Event memory: A theory of memory for laboratory, autobiographical, and fictional events. Psychol. Rev., 122(1), 1–23. doi: http://dx.doi.org/10.1037/a0037907

Schacter, D. L., Addis, D. R., Hassabis, D., Martin, V. C., Spreng, R. N. and Szpunar, K. K. (2012). The future of memory: Remembering, imagining, and the brain. Neuron, 76(4), 677–694. doi: http://dx.doi.org/10.1016/j.neuron.2012.11.001

Scoville, W. B. and Milner, B. (1957). Loss of recent memory after bilateral hippocampal lesions. J. Neurol. Neurosurg. Psychiatry, 20, 11–21. doi: http://dx.doi.org/10.1136/jnnp.20.1.11

Siegler, R. S. (1988). Strategy choice procedures and the development of multiplication skill. J. Exp. Psychol. Gen., 117(3), 258–275. doi: http://dx.doi.org/10.1037/0096-3445.117.3.258

Spiers, H. J., Maguire, E. A. and Burgess, N. (2001). Hippocampal amnesia. Neurocase, 7(5), 357–382. doi: http://dx.doi.org/10.1076/neur.7.5.357.16245

Squire, L. R. (1992). Memory and the hippocampus: A synthesis from findings with rats, monkeys, and humans. Psychol. Rev., 99(2), 195–231. doi: http://dx.doi.org/10.1037/0033-295X.99.2.195

Stoff, D. M. and Eagle, M. N. (1971). The relationship among reported strategies, presentation rate, and verbal ability and their effects on free recall learning. J. Exp. Psychol., 87(3), 423–428. doi: http://dx.doi.org/10.1037/h0030541

Svoboda, E., McKinnon, M. C. and Levine, B. (2006). The functional neuroanatomy of autobiographical memory: A meta-analysis. Neuropsychologia, 44(12), 2189–2208. doi: http://doi.org/10.1016/j.neuropsychologia.2006.05.023

Tulving, E. (1985). Memory and consciousness. Can. Psychol., 26, 1–12. doi: http://dx.doi.org/10.1037/h0080017

Verfaellie, M. and Keane, M. M. (2017). Neuropsychological investigations of human amnesia: Insights into the role of the medial temporal lobes in cognition. J. Int. Neuropsychol. Soc., 23(9-10), 732–740. doi: http://dx.doi.org/10.1017/S1355617717000649

Wechsler, D. (2009). WMS-IV.: Wechsler memory scale-Administration and scoring manual. London, UK: Pearson Assessment.

Woollett, K. and Maguire, E. A. (2010). The effect of navigational expertise on wayfinding in new environments. J. Environ. Psychol., 30(4), 565–573. doi: http://dx.doi.org/10.1016/j.jenvp.2010.03.003

Zeidman, P. and Maguire, E. A. (2016). Anterior hippocampus: the anatomy of perception, imagination and episodic memory. Nat. Rev. Neurosci., 17(3), 173–182. doi: http://dx.doi.org/10.1038/nrn.2015.24

Zola-Morgan, S., Squire, L. R. and Amaral, D. G. (1986). Human amnesia and the medial temporal region: enduring memory impairment following a bilateral lesion limited to field CA1 of the hippocampus. J. Neurosci., 6(10), 2950–2967. doi: http://dx.doi.org/10.1523/JNEUROSCI.06-10-02950.1986

